# Directed evolution of mesophilic HNA polymerases providing insight into DNA polymerase mechanisms

**DOI:** 10.1101/2022.11.08.515583

**Authors:** Paola Handal-Marquez, Leticia L. Torres, Vitor B. Pinheiro

## Abstract

Detailed biochemical characterization of natural and mutant enzymes provides essential clues to understand their mechanisms. There are, however, limits to the throughput of such approaches and they are not without errors. DNA polymerases have benefited from over 50 years of detailed study and remain not fully understood. As such, methods that allow high-throughput interrogation of variants, and viable analysis pipelines to identify relevant variants, become an important tool to accelerate research. Using the DNA polymerase from *B. subtilis* Phi29 bacteriophage as a model, we demonstrate how coupling focused libraries, selection and deep sequencing can be combined to identify variants of interest for characterization. As selection parameters can be controlled, different areas of an enzyme’s mechanism can be explored. Focusing selection on faster HNA (1,5-anhydrohexitol nucleic acid) synthesis, we identified P562del as a variant of interest, enriching significantly between rounds. Characterization confirmed its faster HNA synthesis initiation but lower processivity and fidelity. P562 is a non-conserved residue, unlikely to be selected by more traditional approaches, but its deletion recapitulates knowledge on how Phi29 exonuclease, thumb and TPR2 subdomains regulate polymerase function. Our data further support the hypothesis that Phi29 shows a two-state binding to its template: a fast non-replicative complex that transitions to a replication-competent state.

## Introduction

Information storage and replication is an essential process in all living organisms, and it is carried out by specialist enzymes: template-dependent DNA (or RNA) polymerases. Despite their central role, the ancient origin of those proteins has given evolution time and opportunity to explore a vast array of characteristics and specializations: from high-fidelity highly processive replicative polymerases (e.g. *Bacillus subtilis* Phi29 bacteriophage DNA polymerase (Garmendia et al., 1992) to low-fidelity, low processivity DNA translesion repair enzymes (e.g. the mammalian Pol eta (Waters et al., 2009); from monomeric polymerases to large complexes; from mesophilic to hyperthermophilic enzymes.

Those characteristics and specializations are the result of functional and sequence constraints imposed during the natural evolution of DNA polymerases, and they are not yet fully understood. Replicative DNA-dependent DNA polymerases are often multifunctional enzymes (harboring both polymerase and exonuclease activities) with complex dynamics and operating under multiple, and sometimes overlapping, layers of regulation. For instance, DNA polymerases can often discriminate between their natural substrate (deoxyribose nucleotide triphosphates) and other natural analogues present in the cellular milieu (ribonucleotides, partially phosphorylated nucleotides, and damaged nucleobases), even though the analogues are present at a higher concentration than the substrates (Wang et al., 2012). This is done not only at incorporation in the active site of the enzyme (Tsai & Johnson, 2006; Wang et al., 2012), but also at multiple points during the elongation process (Freund et al., 2022; Khare & Eckert, 2002).

A consequence of their complexity and as a result of their evolution, specific mutations can have different impacts on the function of homologous DNA polymerases. For instance, the *Escherichia coli* bacteriophage T4 has evolved around a high-fidelity DNA polymerase for its successful infection of the bacterial host and even small changes in function, such as loss of 3’→5’ exonuclease activity, can result in a 10-fold reduction of bacteriophage viability (Frey et al., 1993). The same significant drop in viability is not observed in the T4-related RB69 bacteriophage (Bebenek et al., 2001). In another example, mutation of a β hairpin in both T4 and RB69 polymerases leads to loss of fidelity but a similar mutation on the *S. cerevisiae* DNA polymerase δ (also a B-family polymerase) does not. Therefore, the mutational space naturally accessible to each enzyme can differ.

Detailed biochemical and mutational analysis has been very successful at identifying some of the sequence and structural determinants for the different DNA polymerase functions, but they are limited to the models we use to understand polymerases, to how those can be efficiently tested, and to how generalizable they are.

DNA polymerase engineering, whether for DNA applications (e.g. PCR or DNA sequencing) or for the synthesis of xenobiotic nucleic acids (XNAs), has identified multiple new mutations that modulate enzymatic function and further contribute to our understanding of those enzymes (Laos et al., 2014; Pinheiro, 2019). Nonetheless, the gain in functional understanding is not the goal of those efforts, instead being a byproduct of selection and rational design methodologies, and often used as a tool to rationalize the contribution of isolated mutants.

However, those same directed evolution platforms, that rely on high-throughput screening, have the potential to be refocused towards improving our mechanistic understanding of DNA polymerases. While the design of the libraries used in directed evolution creates a boundary for the sequence space that can be explored in a selection campaign, the high-throughput screening used in selection has the power to bypass knowledge gaps: it can operate without the need of a prior hypothesis, and it can overcome previous misinterpretations. The first reduces the likelihood of confirmation biases, while the latter is ideal to challenge established paradigms. In addition, and particularly in the case of polymerases, the use of unnatural substrates that cannot be efficiently incorporated has additional benefits, such as slowing down reaction rates (changing the shape of the sequence space that can be experimentally isolated) and potentially not equally affecting all functional parameters in the polymerases (thus allowing deconvolution of complex interactions).

For example, in engineering a hyperthermophilic DNA polymerase from *Thermococcus gorgonarius* for the synthesis of XNAs, a variant, named 6G12, was isolated for its ability to efficiently synthesize HNA (1,5-anhydrohexitol nucleic acids) (Pinheiro et al., 2012). It harbored a total of 18 mutations: four introduced to facilitate engineering (e.g. disabling its exonuclease activity), and 14 isolated from selection. At the time, we rationalized that reshaping of the enzyme thumb domain was needed to accommodate the distorted hybrid nascent HNA-DNA helix. Nonetheless, later efforts to engineer a mesophilic XNA polymerase from the *B. subtilis* bacteriophage Phi29, determined that reducing (N62D mutation) or disabling (D12A) the exonuclease activity of the polymerase was already sufficient to enable HNA synthesis – albeit substantially less efficient than its DNA activity. An updated hypothesis is, therefore, that stability of the polymerase XNA/DNA duplex is the driving parameter for efficient XNA synthesis, as observed for a DNA polymerase engineered for processive RNA synthesis (Cozens et al., 2012).

Phi29 DNA polymerase is known to be a highly processive enzyme (processivity defined as number of nucleotides incorporated by a single polymerase in a single DNA-binding event), capable of up to 70 kb synthesis per binding event (Lieberman et al., 2010). Phi29 processivity is closely associated with the terminal protein region 2 (TRP2) subdomain, which forms a structure akin to a sliding clamp around template and nascent DNA, and which harbours multiple contacts with the thumb (along residues V559-V566) and exonuclease (along residues T15-V24 and F65-N72) subdomains (see **Figure 1A**).

**Figure 1.**
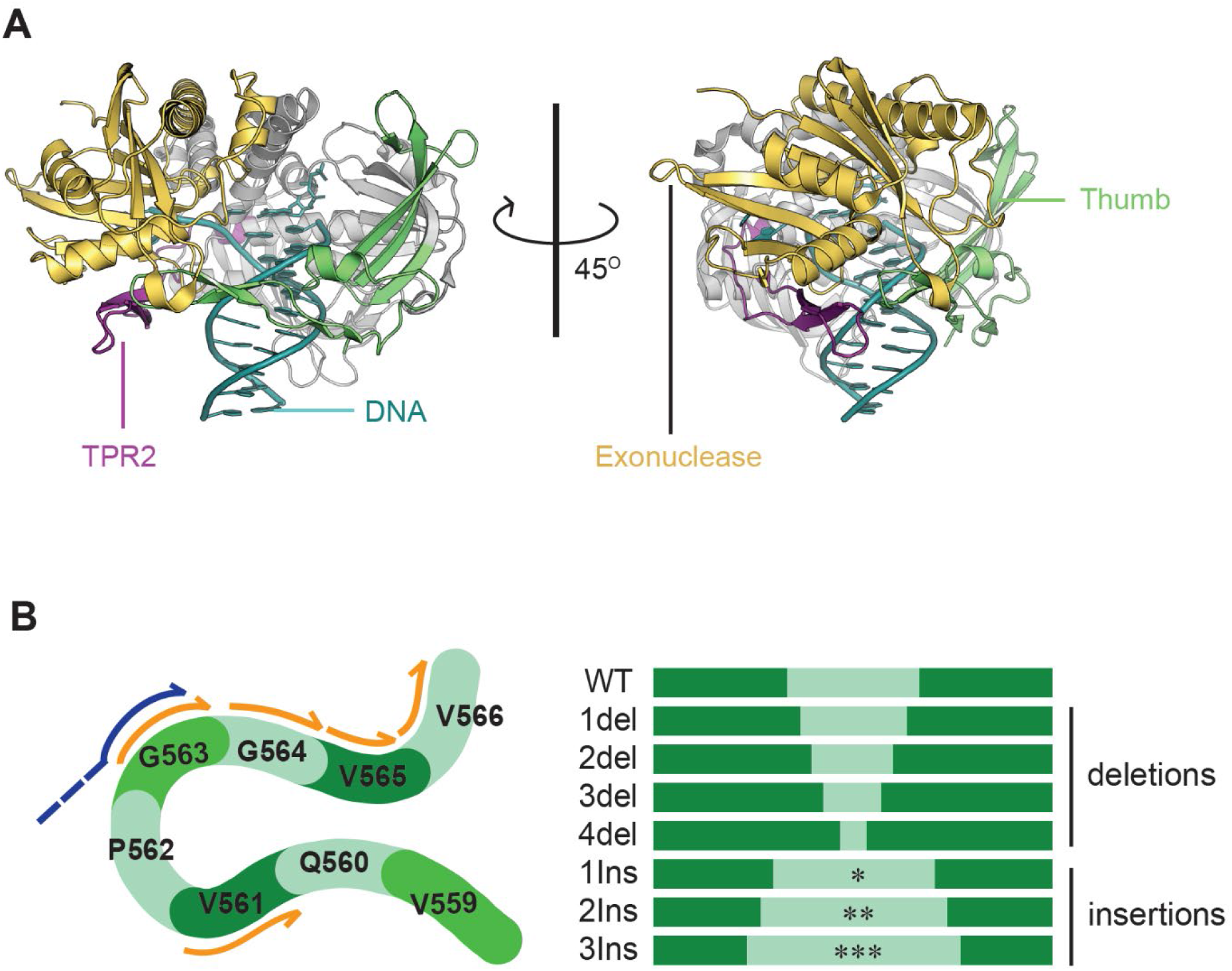
Structure of the closed ternary complex of Phi29 DNA Polymerase and InDel library design. (**A**) There domains in the phi29 DNA polymerase (PDB: 2PYJ) are involved in creating a clamp around the nascent DNA duplex: the polymerase thumb (green), the exonuclease domain (yellow) and the TPR2 domain (purple). The DNA duplex is shown in cyan. (B) InDel mutagenesis of the thumb loop through inverse PCR (iPCR). A single reverse primer in combination with forward primers harbouring 1-3 NNS codons or with priming sites that skipped 1-4 codons were used to generate focused libraries or deletion mutants respectively. The same approach was used to target the TPR2 and exonuclease loops.

Deletion of the TRP2 subdomain, does not affect the ratio between exonuclease and polymerase activities of the polymerase but significantly decreases polymerase affinity for DNA and its processivity (Rodriguez et al., 2005). This is analogous to what is observed in the T7 bacteriophage DNA polymerase (gp5), where high processivity is only observed when a clamp-like structure is formed around the DNA by the binding of the thioredoxin processivity factor (Tran et al., 2012).

The close contacts between TRP2, thumb and exonuclease can, in principle, affect other polymerase processes, including exonuclease activity (given the packing against an alpha helix harbouring one of the catalytic exonuclease residues – D66), exonuclease sampling (DNA switching between polymerase and exonuclease catalytic sites, by modulating how exonuclease and thumb interact), strand displacing activity (through the interaction TRP2 with template DNA) and DNA binding.

This three-way junction is therefore also a relevant region for the engineering of efficient HNA polymerases based on the Phi29 DNA polymerase chassis, as the optimisation of the sequence length and composition in these loops could potentially create new molecular interactions that stabilize protein binding to the nascent heteroduplex or avoid steric clashes that could be contributing to the lower processivity observed during HNA synthesis (Handal Marquez, 2019). If any of the other processes described above can be affected by changes in this region, that could also lead to improvements in HNA synthesis.

The potential role of enzyme processivity on HNA synthesis, together with the biochemical evidence of how TRP2 is involved in modulating it, establish a tractable system for investigating whether selection platforms can be used to gain biological insights into complex enzymatic mechanisms. Libraries can be designed with minimal assumptions, selection conditions can be tailored towards investigating specific parameters, and deep sequencing analysis of the selection output rapidly identifies variants of interest: enzymes enriched by the selection conditions.

Here, we demonstrate how this system can be assembled and we use it to identify a single residue in the DNA polymerase thumb (P562) implicated in Phi29 DNA polymerase processivity. We characterise the isolated enzyme to demonstrate the impact of the mutation on DNA affinity, enzyme processivity, fidelity and strand displacement activities. We confirm that Phi29 DNA polymerase needs not only to bind DNA but also to switch to a stable conformation to achieve highly processive DNA synthesis.

## New Approaches

We demonstrate that high-throughput selection can be harnessed for high-throughput biochemistry, allowing novel mechanistic insights in enzyme function. The approach relies on an integrated approach to library design, selection and data analysis (**Figure 2**), so that relevant enzyme variants can be identified and further characterised. The approach is scalable but hinges on the effectiveness and specificity of the selection strategy being implemented.

**Figure 2:**
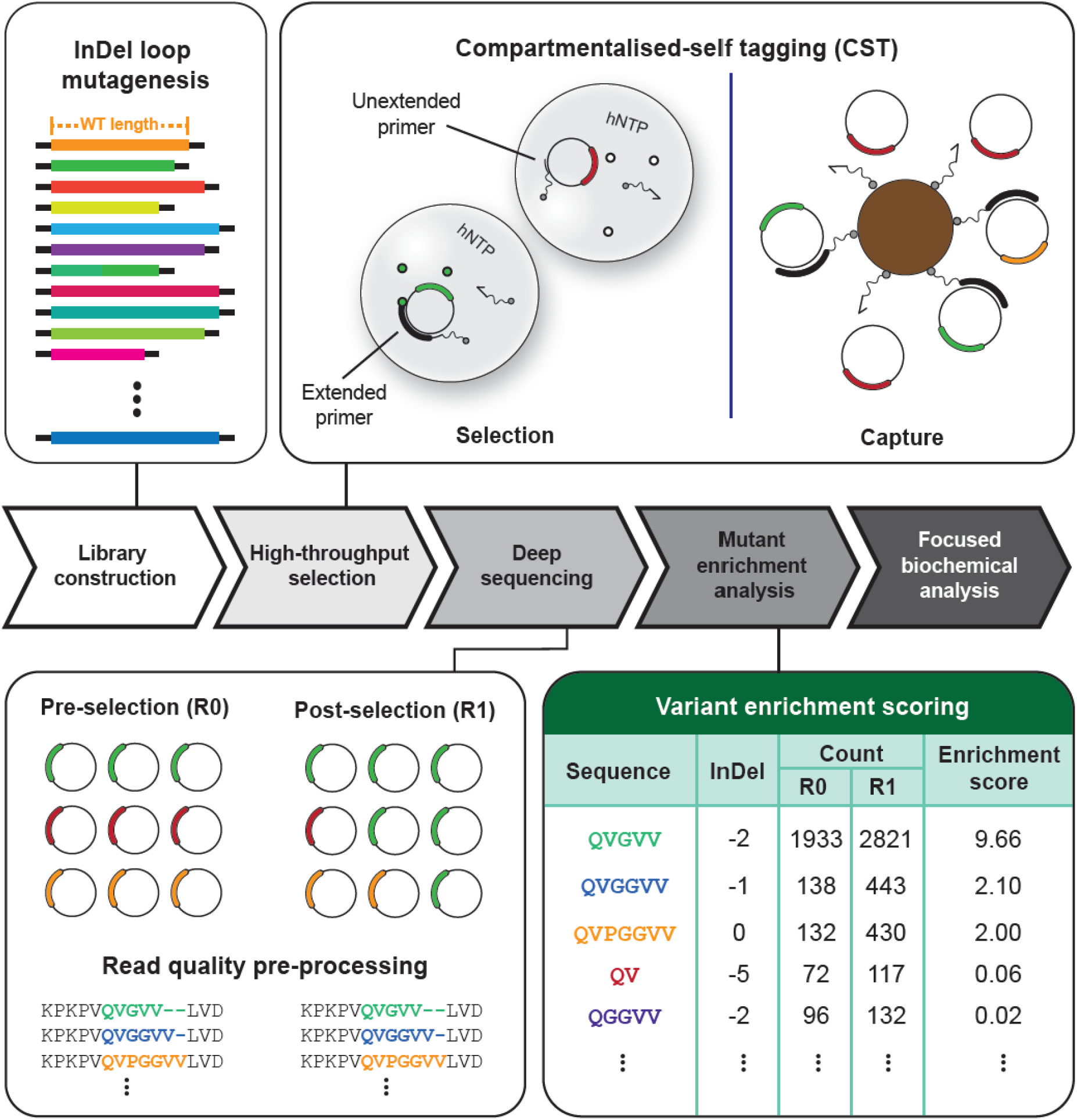
Mesophilic XNA polymerase directed evolution pipeline. A library is initially constructed – represented (top left panel) by different color/lengths compared to the wild-type. The library is then subjected to a round(s) of selection through compartmentalized self-tagging (top right panel). The library pre- and post-selection is sequenced through NGS (bottom left panel). The proportion of highly active variants (e.g., green) should increase whereas that of variants with lower activity (e.g., orange, and red) should decrease. NGS data is filtered for quality. Pre-processed data is then analyzed (bottom right). The frequency of each mutant corresponds to its abundance in the library pre- and post-selection. The enrichment score is used to evaluate the increase or decrease in abundance of specific mutants within the population and serves as starting point for further analysis.

## Results

### Library design: focused libraries sampling beyond biochemical knowledge

There are four interconnected steps that are crucial for harnessing selection as a tool to investigate biology: library design, selection methodology, selection conditions and downstream data analysis.

The wide range of available DNA library assembly strategies allows length and compositional variation to be introduced, randomly or in a targeted manner, across a gene of interest (Chica et al., 2005; Lutz, 2011), such that technical bottlenecks in library generation are no longer relevant. Instead, design is constrained by the number of variants that can be generated and sampled (linking design to selection methodology), considering the likely destabilizing effect of a high mutational burden to the protein being engineered.

Although there are multiple possible contacts between TRP2, thumb and exonuclease subdomains, we hypothesised that contact between the three loops (TRP2: L406-L412, thumb: V559-V566, and exonuclease T15-V24) is likely functional, capable of providing the dynamic response needed for polymerase-exonuclease balance and enzyme processivity. Only limited diversity can be introduced in the 29 residues, given the rapid rise in the number of variants and the poor stability of the Phi29 DNA polymerase (half-life of 37 ± 4.5 min at 40°C, when in the presence of stabilizing DNA template - (Povilaitis et al., 2016).

Starting from a previously reported thermostabilized Phi29 DNA polymerase (Povilaitis et al., 2016) (t_1/2_ > 16 h at 40°C in the presence of DNA), harboring the additional D12A mutation (which disables exonuclease activity) to allow HNA synthesis, we opted to concomitantly explore loop length in the three loops. Firstly, because insertions and deletions (indels) tend to be least disruptive to protein stability when they occur in loops. Secondly, we hypothesized that loop distortions, due to indels, would be a more sensitive tool to study loop-loop interaction than patterns of substitution, while also minimizing library diversity. That is further supported by sequence alignment with Phi29 DNA polymerase homologues (**Figure S1**), showing that these regions can accept indels.

Secondly, given that the HNA-DNA heteroduplex is expected to be wider than a DNA duplex, we hypothesized that an insertion could lead to stabilizing interactions between the three subdomains, disrupted by the wider substrate, being re-established – as reported for the engineering of the *Chlorella* virus DNA ligase towards an XNA ligase (Vanmeert et al., 2019).

The tips of the loops (residues D19, N409 and P562) were used as seed in design, with up to three insertions being introduced upstream of those seeds, or up to four deletions targeting the seed and downstream residues, as shown in **Figure 1B**. Exonuclease, TRP2 and thumb libraries had each a theoretical maximum diversity of approximately 8425 variants – well within the sampling range of the chosen selection platform (approximately 2 × 10^7^ per experiment) and analysis pipeline (between 5.0 and 7.0 × 10^4^ of sequences per run).

Although guided by the polymerase structures and phylogeny, the libraries are agnostic to the biochemical knowledge that TRP2 (Rodriguez et al., 2005; Rodríguez et al., 2009) and the Phi29 DNA polymerase C-terminus (Kamtekar et al., 2004; Truniger et al., 2004) are crucial for processivity, with a single previous report targeting one of those residues (Rodríguez et al., 2009).

### Using enrichment to identify relevant mutations in selection

High-throughput selections rely on systems that link the phenotype being sought with efficient recovery of the responsible genotype, so that at each round of selection, the genotype of active enzymes becomes more frequent.

A variety of selection methodologies have been demonstrated for the selection of DNA and XNA polymerases, including *in vivo* (Patel & Loeb, 2000), phage display (Xia et al., 2002), and emulsion-based methods such as CSR (compartmentalised self-replication) (Ghadessy et al., 2001), CST (compartmentalised self-tagging) (Pinheiro et al., 2014), and CBL (compartmentalised bead-labelling) (Houlihan et al., 2020). Technical characteristics of each method shape the range of polymerase activities that can be selected and what activities can emerge as selection parasites (active enzymes that co-enrich with the desired activity).

CST relies on polymerases capable of stabilizing the binding of the selection primer against their own gene through XNA synthesis (**Figure 2**). Given the diminishing return per XNA incorporation to the overall primer stabilization (i.e. analogous to oligonucleotide length effects on primer melting temperature), the pressure of CST selection is highest on the first few incorporations, with CST selections working akin to a high-pass filter (i.e. selection is unable to discriminate between polymerases that can synthesise beyond a certain number of incorporations). Expected parasites in this CST selection include polymerases that can make use of the low concentrations of natural cellular dNTPs still present, polymerases that express to significantly higher levels or that are more thermostable.

For the selection of HNA polymerases with higher processivity, extension time is a key parameter in selection, since shorter extension times are likely to benefit highly processive enzymes as well as polymerases that can start XNA synthesis faster. The latter was not expected to be a significant possibility, considering Phi29 high affinity for DNA (K_D_ = 1μM in conditions close to the selection conditions (Lázaro et al., 1995)) and in view of our experience engineering thermostable XNA polymerases.

A single round of selection for HNA synthesis was carried out for each library, followed by deep sequencing of naïve and selected libraries for analysis. Frequency counting post-selection can be a suitable route to identify best candidates in circumstances where libraries are not biased, for selection methodologies that have very low background (i.e. recovered genotypes that are randomly recovered) and for late-stage selection campaigns (i.e. libraries that have already been carried through multiple selection rounds and that are populated by few variants) (Wrenbeck et al., 2017). As our libraries were pooled (by site across all designed lengths) during assembly, we expected them to show significant coverage biases. In addition, emulsion-based polymerase selection platforms show significant background due to PCR amplification post-selection. Therefore, we opted to focus on enrichment, rather than simply frequency, as a measure of selection.

Sequencing errors and suboptimal sampling of the libraries also contribute to the complexity of traditional analyses, with most simple enrichment estimators unable to cope with sequences that appear only in the selected pool. Treating the observation of individual sequence variants as independent Poisson distributions permits pre- and post-selection populations to be tested for statistically significant differences, via an E-test (Krishnamoorthy & Thomson, 2004). However, multiple testing across all variants rapidly limits the usefulness of the approach.

Instead, we filtered the datasets to focus on variants that were present in both datasets and that increased in frequency between pre- and post-selection. In addition, we scored them not only on their change in frequency but also on their average number of observations, to minimise false positive detection by concentrating analysis on better sampled variants. Significantly enriched variants are shown in **Table 1**.

**Table 1.**
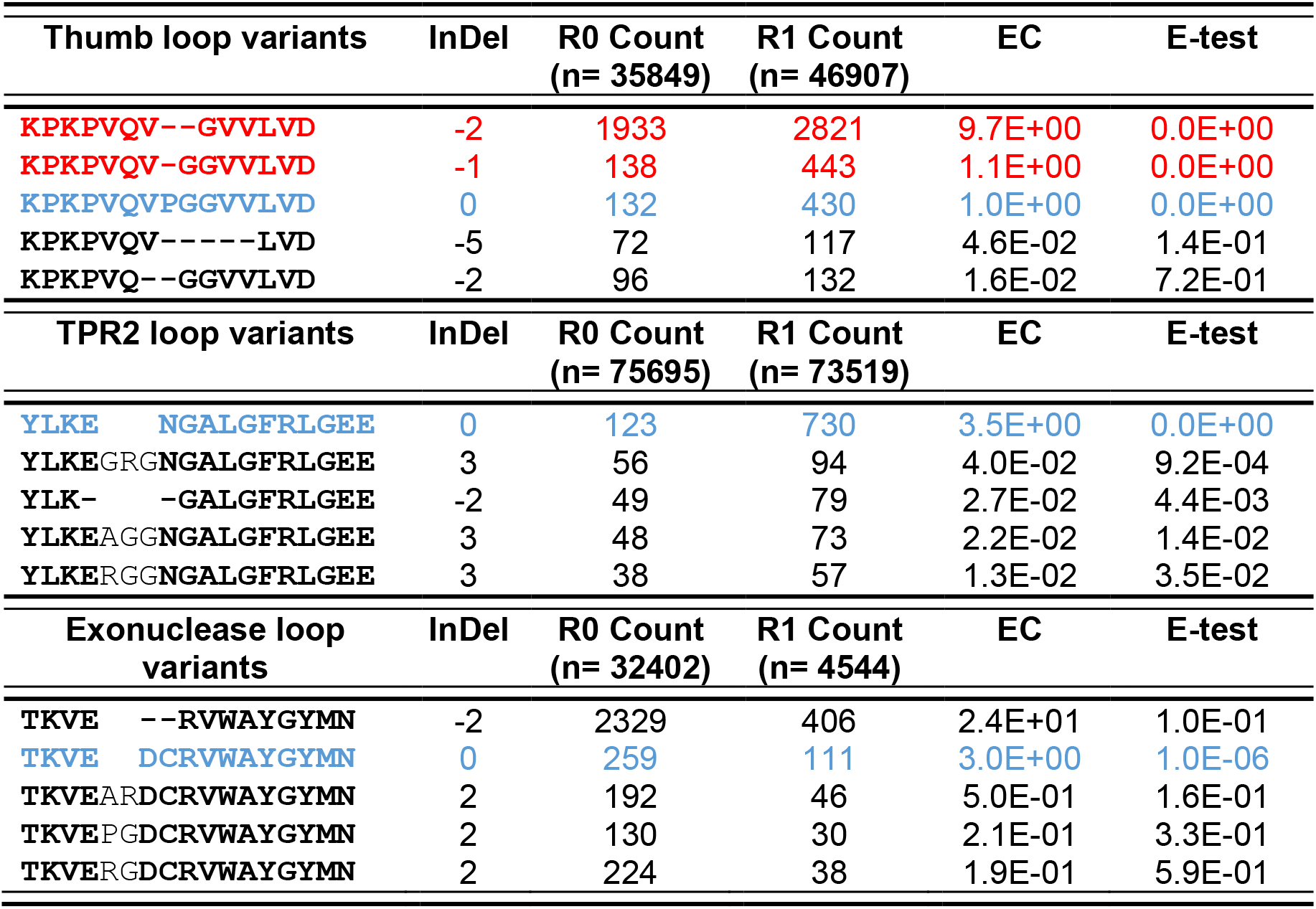
Most enriched variants isolated from individual libraries. Complex enrichment (EC) scores and statistical significance for the top 5 variants from each of the phi29 DNAP Exonuclease, TRP2, and thumb InDel loop libraries. The EC is calculated as the difference in frequency between rounds times the average number of counts. Wild-type sequences and their corresponding scores are shown in blue. Sequences with higher EC than the wild-type and significantly enriched (E-test p-value <0.05) are shown in red. Deletions are shown as hyphens and insertions are shown in not in bold.

Out of the identified mutants, P562del (Phi29 DNAP M8R, D12A, V51A, M97T, G197D, E221K, Q497P, K512E, F526L, ΔP562) and PG563del (Phi29 DNAP M8R, D12A, V51A, M97T, G197D, E221K, Q497P, K512E, F526L, ΔP562-G563) variants were notable, having enriched better than the parental enzyme (D12A-THR: Phi29 DNAP M8R, D12A, V51A, M97T, G197D, E221K, Q497P, K512E, F526L) in selection. In view that deletion of P562 alone was sufficient for the increased enrichment, we chose to focus on this variant for further characterisation.

### Deletion of P562 reduces processivity and fidelity

Although kinetic parameters of incorporation are often used to describe DNA polymerase activity and fidelity (when carried out with non-cognate substrates), they often do not consider effects that emerge during elongation, which are known to be significant for XNA synthesis (Lutz et al., 1999; Xia et al., 2002). For instance, hexitol nucleotide triphosphate (hNTP) incorporation kinetics is comparable to dNTP kinetics (Vastmans et al., 2000) in the B-family Vent DNA polymerase, yet Vent cannot incorporate more than 7 hNTPs in a row.

Therefore, initial characterisation focused on confirming the increased activity of P562del for HNA synthesis by primer extension using concentration-normalised purified enzymes – confirming the results of selection and analysis. In short reactions (60 min extension), as shown in **Figure 3**, P562del indeed outperforms the starting D12A-THR polymerase, synthesising longer HNAs (~37 incorporations vs. ~18 from D12A-THR) and more efficiently (less visible unextended primer). Longer incubations (**Figure S2A**), however, allow D12A-THR to match and eventually outperform P562del – differences that cannot be seen with the natural nucleoside triphosphates. Notably, like the previously reported ΔTPR2 mutant (Rodriguez et al., 2005), HNA synthesis by the P562del is distributive while D12A-THR is processive (with few visible accumulated intermediates between primer and full-length extension). Primer extension using alternative templates ruled out possible sequence-specific effects (**Figure S2B**).

**Figure 3.**
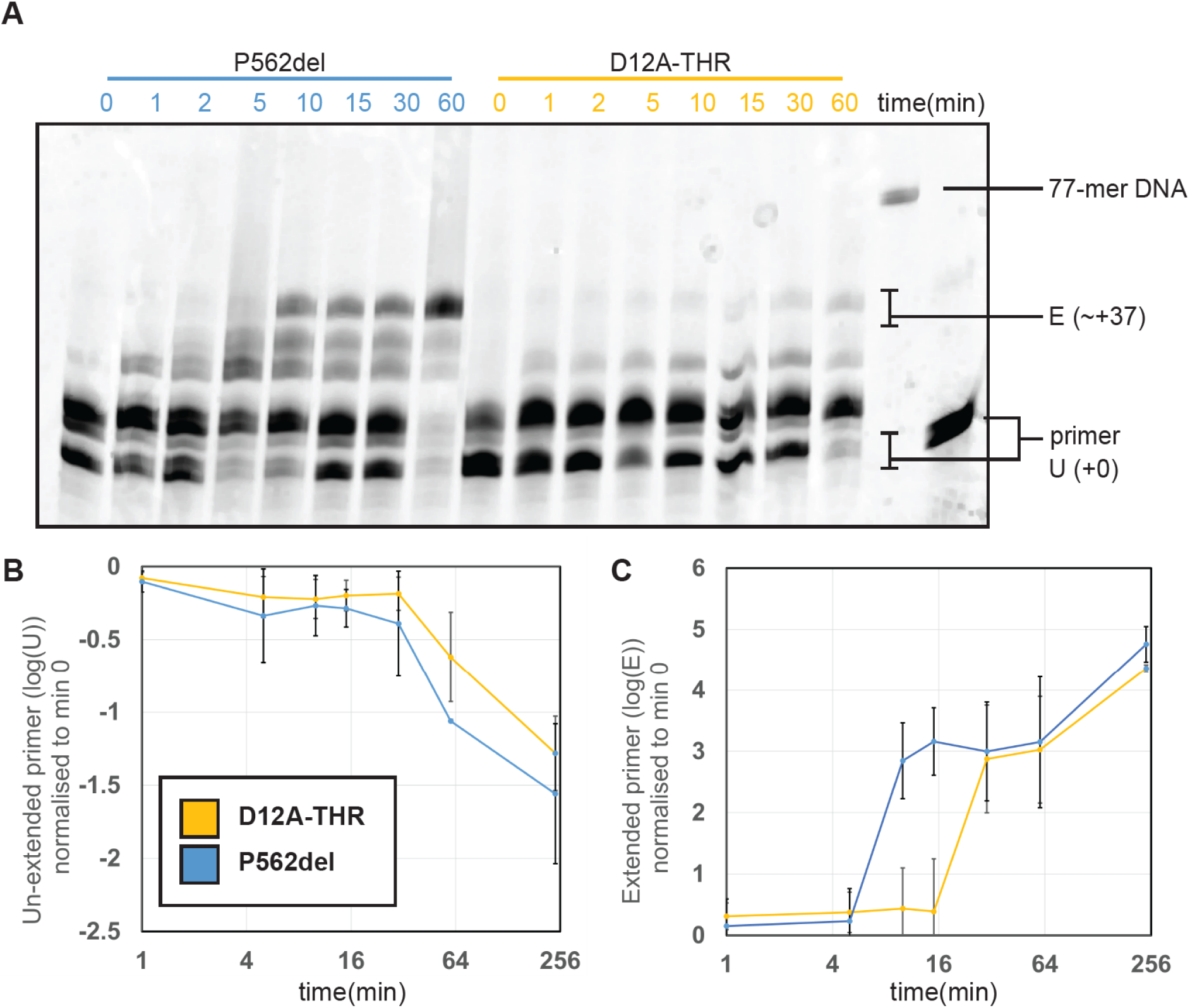
P562 deletion in phi29 DNAP enables faster initiation of HNA synthesis. Primer extension assays for HNA synthesis were carried out by incubating 10 pmol of single stranded DNA template pre-annealed to 1 pmol of fluorescently labelled DNA primer with 12 nM D12A-THR or P562del phi29 DNAP over different amounts of time. (**A**) HNA primer extension products synthesized by D12A-THR and P562 mutants separated by denaturing PAGE. Unextended primer (U) and extension products of ~37 incorporations (E) are highlighted. (**B**) Average depletion of unextended primer (U) by D12A-THR (orange) and P562del (blue) normalized to the earliest time point (0 min). (**C**) Average extended products with ~37 incorporations (E) by D12A-THR and P562del normalized to the earliest time point (0 min). P562del shows a faster start to its hNTP incorporations than D12A-THR.

Multiple polymerase mechanisms could account for the “early burst” phenotype observed. For instance, changes in binding affinity to the template (which begins as dsDNA but changes into an HNA/DNA heteroduplex as the polymerase extend the primer), changes in processivity (by increasing the dissociation rate of the enzyme from the template), or a drop in fidelity (misincorporations stall the polymerase and can also lead to more frequent sampling of the editing conformations). We therefore focused characterization on assays that could give some insight into those processes, always comparing P562del, to the parental enzymes, with (Exo+THR: Phi29 DNAP M8R, V51A, M97T, G197D, E221K, Q497P, K512E, F526L) or without (D12A-THR) active exonuclease function.

Electrophoretic mobility shift assays (EMSAs) showed a significant difference between P562del and D12A-THR (**Figure 4)**. The parental enzyme, as previously reported, shows complexes with two different mobilities: an unstable intermediate that transitions to a stable replication-competent complex (Rodriguez et al., 2005). The deletion of P562 changes that process, with both complexes co-existing even after long incubations, suggesting that the stability of the replication-competent complex is reduced.

**Figure 4.**
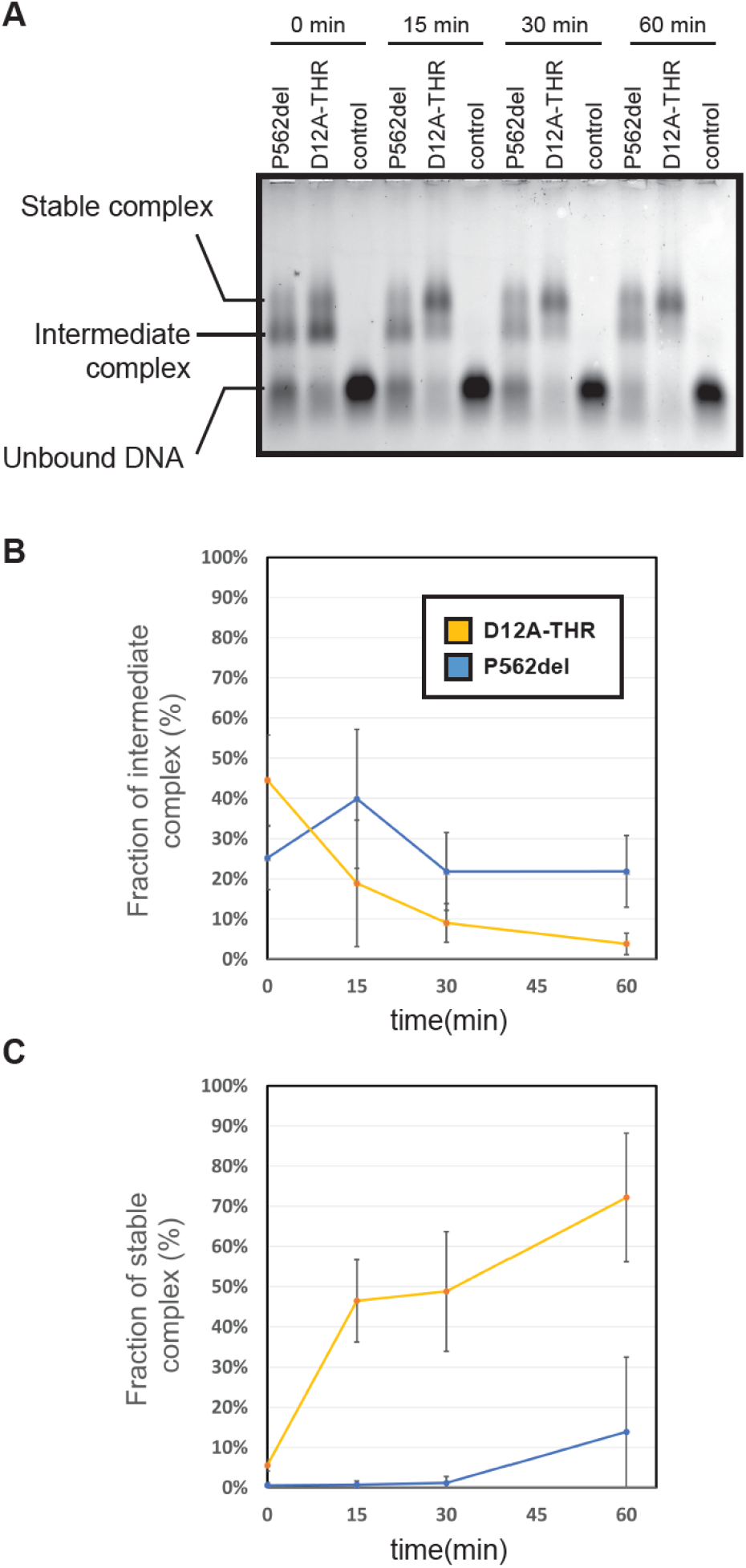
Phi29 DNAP P562del reduced DNA binding capacity. For the EMSA, 60 pM commercial phi29 DNAP (control), D12A-THR or P562del were allowed to bind a fluorescently labelled primer pre-annealed to a ssDNA template (**A**) Reactions with commercial phi29 DNAP show no binding, probably as a result of storage buffer components interfering with quantification. (**B**) Fraction of intermediate Pol-DNA complex, and (**C**) the fraction of stable Pol-DNA complex by D12A-THR (orange) and P562del (blue) over time.

Rolling circle amplification (RCA) confirmed the detrimental impact of D12A (Bernad et al., 1989) and showed that deletion of P562 results in an enzyme unable to perform RCA (**Figure 5A**). Average extension lengths appear shorter than D12A-THR and no full-length product could be observed, supporting that P562 may be an important contact between thumb and TPR2 subdomains, and essential for TPR2’s role in strand displacement and high processivity. In the context of the previous primer extension and EMSA results, this supports a role for P562 in the dissociation rate of the polymerase from its DNA template.

**Figure 5.**
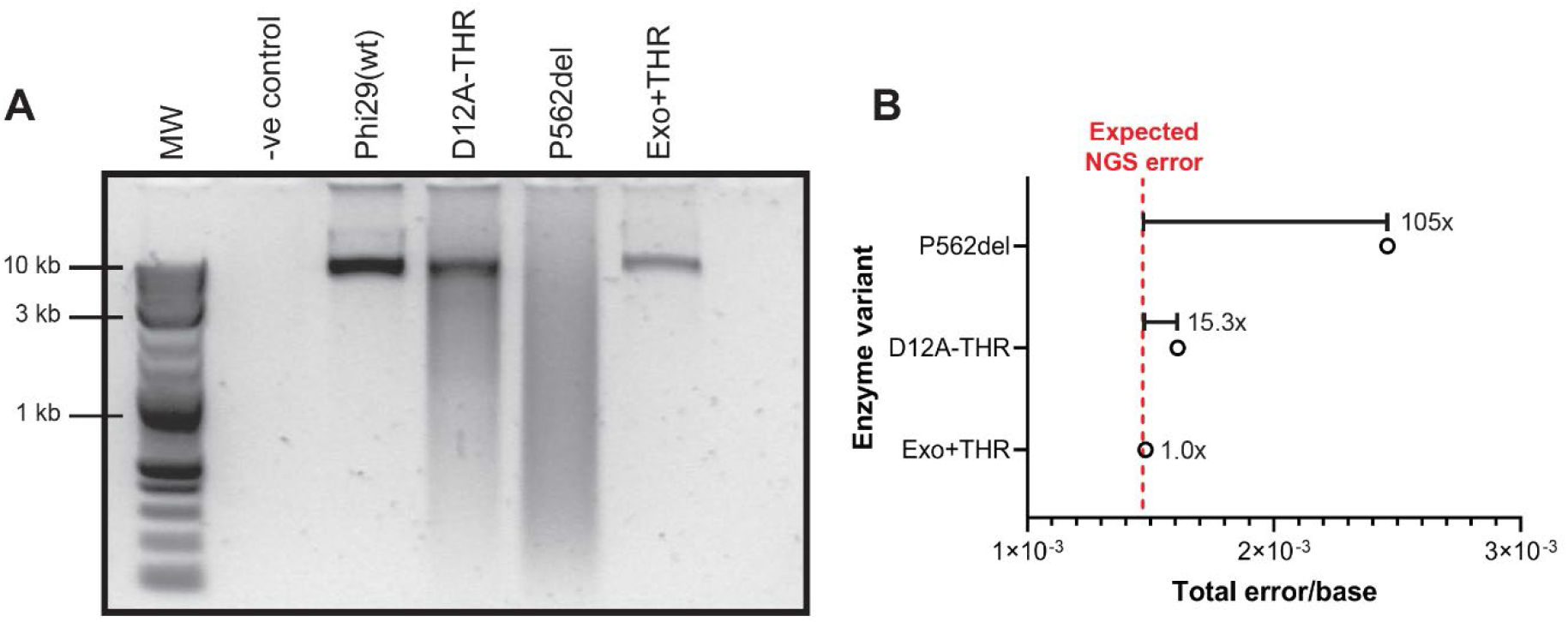
Impaired rolling circle amplification (RCA) activity and reduced fidelity Phi29 DNAP P562del. (**A**) The RCA assay used 10 ng of plasmid template and 3 nM of each enzyme in a 3-hour reaction. A sample without enzyme (-ve control) was used to confirm the polymerase-dependent amplification. Restriction of the RCA yields full-length products. (**B**) Relative fidelity of the D12A-THR and p562del mutants to Exo+THR. The corrected total error rates from **Table 2** of each mutant were divided by that of the Exo+THR, resulting in the relative fidelity scores.

DNA synthesis fidelity was investigated using an adaptation of our previously reported XNA fidelity assay, based on primer extension assays (**Figure S3A**). Given the very low error rates shown by wild-type Phi29 DNA polymerase (in the range between 3×10^−6^ (Nelson et al., 2002) and 9.5×10^−6^ (Paez et al., 2004) errors per base), errors introduced by PCR and deep sequencing steps significantly affect the data obtained. Deep sequencing errors alone are reported around 4.73 ×10^−3^ for MiSeq Illumina platforms(Stoler & Nekrutenko, 2021).

We opted to use Exo+THR fidelity data to normalize for PCR and deep sequencing errors in the experiment, thus establishing a baseline against which the impact of D12A and P562 could be measured. As expected, disabling the exonuclease activity (D12A-THR) increased the rate of substitutions by approximately an order of magnitude, while showing marginal increases in deletions and insertions. Removal of P562 had a more pronounced effect, with substitution rates approximately 100-fold higher than Exo-THR and significant increases in deletion (7-fold) and insertion (6-fold) frequencies (**Figure 5B** and **Table 2**).

**Table 2.** Quantification of insertions, deletions, and substitutions rates (errors/bp) of phi29 DNAP mutants. ^a^Raw error rates that do not consider PCR and NGS error rates. ^b^R: PCR and Error estimate calculated from the number of errors expected to be introduced by the commercial phi29 DNAP (using the published 9.5×10^−6^ error rate (Paez et al., 2004)) and subtracting this figure from the observed number of errors introduced by Exo+THR. The remainder was used to calculate the expected PCR and NGS error estimate for the total bases sequenced of the D12A-THR and p562del mutants.

| Mutant | Substitution rate<sup>a</sup> | Deletion rate<sup>a</sup> | Insertion rate<sup>a</sup> | Total error rate | R<sup>b</sup> | Corrected total error rate | Total bases |
| --- | --- | --- | --- | --- | --- | --- | --- |
| Exo+THR | $1.4 \times 10^{-3}$ | $9.0 \times 10^{-5}$ | $4.0 \times 10^{-6}$ | $1.5 \times 10^{-3}$ | $2.6 \times 10^3$ | $9.5 \times 10^{-6}$ | 1,754,723 |
| D12A-THR | $1.4 \times 10^{-3}$ | $1.7 \times 10^{-4}$ | $6.1 \times 10^{-6}$ | $1.6 \times 10^{-3}$ | $2.7 \times 10^3$ | $1.5 \times 10^{-4}$ | 1,808,769 |
| P562del | $1.8 \times 10^{-3}$ | $6.1 \times 10^{-4}$ | $2.3 \times 10^{-5}$ | $2.5 \times 10^{-3}$ | $2.7 \times 10^3$ | $9.9 \times 10^{-4}$ | 1,834,463 |

Detailed analysis of error identity and position showed context-dependent mutational hot spots for the three tested enzymes but particularly salient for P562del (**Figure S4-S5**). P562del also showed elevated levels of transition substitutions (26.8%) compared to D12A-THR (21.2%) – shown in **Figure S3C**.

The phenotype shown by P562del bear similarities to the previously reported ΔTPR2, in that the variants form a less stable complex with their template, resulting in lower processivity and loss of strand displacement activity. That, however, is not sufficient to explain P562del enrichment in selection and its increased initial rate of HNA synthesis. Given the previous report of P562 interaction with TPR2 (Rodriguez et al., 2005), we postulate that P562 deletion removes a critical contact involved in regulating Phi29 polymerase binding to its template, making the process more dynamic – faster at assembling replication-competent complexes (explaining selection and primer extension), and faster at dissociating from DNA (explaining lower DNA binding stability and poor RCA performance). Dissociation from template becomes even more likely once it becomes an HNA/DNA heteroduplex, as it has been observed for other polymerases (Vastmans et al., 2000), explaining the “early burst” phenotype observed.

## Discussion

DNA polymerases are essential for life and, logically, they have been extensively studied in the last 60 years. In addition to biochemical characterisation of natural enzymes, mutants – whether selected or targeted – have also significantly contributed to our current detailed mechanistic understanding of this family of enzymes (Pinheiro, 2019).

Nonetheless, that understanding remains incomplete for polymerases. For instance, we still have no mechanistic understanding of how some natural DNA-dependent DNA polymerases display reverse transcriptase activity – such as the DNAP from *Thermus thermophilus* for RNA (Myers & Gelfand, 1991), or the DNAP from *Geobacillus stearothermophilus* for some XNAs (Jackson et al., 2019). We do not know how to explain the different impact of a given modification between different polymerase chassis (Nikoomanzar et al., 2019), and we still compare enzyme activities at standardized conditions, rather than at their individual reaction optimum (Nikoomanzar et al., 2017). Such lack of understanding also explains the current generation of XNA polymerases, which lag severely behind natural DNA polymerases (using natural substrates) on processivity, fidelity and catalytic rates.

Numerous individual residues have been implicated in function, but they cannot act in isolation. How the different residues interact as a system to enable a polymerase to function remains elusive, with only a handful of examples demonstrating that such systems exist (Li et al., 2010).

For less well-characterised enzymes, the prospect of biochemically characterising panels of mutants becomes increasingly difficult, and only accessible by technically challenging high-throughput screening platforms (Gielen et al., 2016). Still, selection platforms have the potential to explore even higher throughputs (> 10^7^ variants) and at a lower technical entry barrier (e.g. emulsion or *in vivo* platforms).

Selection platforms are at the core of directed evolution, but here the emphasis is changed: selection conditions are chosen not to enhance certain feature of the enzyme but to recover functional enzymes that perform best at the selected conditions. Consequently, there is greater need for robust statistical analysis of selection outputs, and for a more systematic exploration of how reaction conditions influence selection – areas that increasingly being considered in directed evolution.

We implement an E-test to compare how individual sequences enrich between rounds, approximating individual sequence frequencies to Poisson distribution parameters. It assumes a degree of robustness in selection (so that frequencies can be good population estimates) that is only justifiable for heavily oversampled libraries – here, designed libraries were under 10^4^ variants, transformations recovered in excess of 10^6^ CFU and CST selections are carried out with over 10^7^ cells, fulfilling those criteria. Nonetheless, a natural progression would be to include stochastic measures that assume incomplete sampling, as commonly implemented for aptamer selection (Hoinka et al., 2015), and to extend analysis to include interacting residues (Nikoomanzar et al., 2019; Salinas & Ranganathan, 2018).

Another crucial aspect of this approach is the library design. Substitutions are traditionally used to sample sequence space around wild-type enzymes, but they do not necessarily abolish backbone interactions. The P562 residue is not conserved among homologous Phi29-like enzymes, and a P562C mutation has been shown to have little functional impact in reducing conditions (Rodríguez et al., 2009), suggesting that its interaction with TPR2 and exonuclease domains may be the result of backbone interactions.

Indels, while abundant in nature, are usually less explored as a tool for protein engineering (Tizei et al., 2021; Tóth-Petróczy & Tawfik, 2013), because of their expected impact on protein structure and increased complexity in analysis.

While this work represents a proof-of-principle that selection can be used to gain biological insight into the workings of a well-characterised enzyme, it can be readily implemented to other enzymes and at larger scales, towards reconstructing the functional networks that define a protein function.

## MATERIALS AND METHODS

### Library design, selection, NGS and analysis leading to the identification of P562del

The exonuclease, TPR2 and thumb loop InDel libraries were generated through inverse PCR (iPCR) on the pET23-P2-D12A-THR (**Table S1**) with phosphorylated InDel mutagenic primers (Table S2) as previously described (Handal Marquez, 2019) and summarized in **Figure 1B**. Each InDel library was subjected to rounds of selection through CST (Pinheiro et al., 2014) adapted for the mesophilic phi29 DNAP as previously described (Handal Marquez, 2019). Briefly, 1×10^8^ cells from each library, post-induction (3 h at 30°C), were resuspended in 100μl of activity reaction mix, composed of 30 pmol of a short biotinylated oligo (CST_04(7)exoR, **Table S2**), 200μM of hNTPs, 1× Phi29 reaction buffer (NEB), 1× Bovine Serum Albumin (BSA, Sigma Aldrich), 1 M betaine (Thermo Scientific), 2μL NotI (NEB), 1 mg/ml lysozyme (Thermo Scientific) and 5μg/ml polymyxin (Sigma-Aldrich) in 100μL molecular grade water. Resuspended cells were emulsified in water-in-oil emulsions and subjected to freezing-thawing cycles for cell lysis and plasmid denaturation. HNA synthesis was then carried out for 10 min at 30°C and products were captured, post-emulsion disruption, via biotin-streptavidin pulldown using paramagnetic beads. Captured products were amplified through PCR using KOD Xtreme Hot Start DNA Polymerase (EMD Millipore), cloned into the pET23-P2-D12A-THR expression backbone, and transformed into fresh electrocompetent *E. coli* T7 Express cells (NEB). The libraries pre- and post-selection were then grown for 3 h at 37°C. Plasmids were then extracted and used as template in ~400 bp NGS amplicon generation PCR reactions using KOD Xtreme Hot Start DNA Polymerase (EMD Millipore). The amplicons were gel-extracted using the Monarch DNA Gel Extraction kits (NEB). NGS-based amplicon sequencing was carried out for all libraries at Genewiz UK Ltd using the Amplicon-EZ service. Sequencing data, which can be found in the NCBI SRA database (BioProject: PRJNA883233), was processed using the *NGS_preprocessing.ga* script (**Supplementary Information S2**) in the Galaxy (Afgan et al., 2016) public server (usegalaxy.org). InDel frequencies before and after selection, enrichment complex scores and the E-test for comparing two Poisson means (Krishnamoorthy & Thomson, 2004) were calculated using the *InDel_Quantification.jl* script (**Supplementary Information S3**). The calculated scores can be found in the *InDel_EC.xlsx* file (**Supplementary Information S4**).

### Generation of phi29 mutants

The P562del and Exo+THR mutants were generated on the pET23-P2-D12A-THR (Table S1) background through inverse Polymerase Chain Reactions (iPCR) using Q5^®^ High-Fidelity DNA Polymerase (NEB) following the manufacturer’s instructions in 25 μl reactions with primers p2_thumb_loop_R/DEL1 (**Table S2**) and iPCR_P2_Exo+_F1/R1 (**Table S2**), respectively. Amplified products were treated with 0.4U/μl DpnI for 1 h at 37°C prior to the PCR purification with the GeneJET PCR Purification Kit (Thermo Scientific). 100 ng of DNA products were blunt-end ligated in 20 μl reactions with 40U/μl T4 DNA ligase in 1× T4 DNA ligase buffer overnight at room temperature. 5 μl of the ligation products were transformed in NEB^®^ 5-alpha competent *E. coli* cells following the recommended High Efficiency Transformation Protocol (C2987, New England Biolabs) described by the commercial strain provider. Successful cloning was confirmed through Sanger Sequencing in all cases.

### Expression, purification and protein quantification of mutants

The pET23 plasmid encoding the phi29 DNA polymerase mutants were transformed into *E. coli* T7 Express cells (NEB). Isolated transformants were grown at 37°C until an OD_600_ of 0.8 was reached. Protein expression was induced by adding IPTG to a final concentration of 1 mM and incubating cultures for 4 h at 30°C. Cells were then pelleted through centrifugation at 5000x*g* and the supernatant was discarded. Pellets were frozen for an hour at −20°C and resuspended in 4 mL of B-PER Reagent (Thermo Scientific) per gram of cell pellet with 50 mg/mL lysozyme and 250U/uL Benzoase (Sigma Aldrich NV). Cells were subsequently incubated at room temperature for 15 min. Protease inhibitor (1 mM Pefabloc) and reducing agents (1 mM DTT) were added, and cell debris removed through centrifugation at 5000×*g*. The supernatant was diluted 3-fold in 50 mM Tris-HCl pH 8.0, 1 mM DTT and 1 mM Pefabloc. Expressed protein was purified using the HisPur™ Ni-NTA Resin (Thermo Scientific) using 50 mM Tris-HCl pH 8.0, 1 M NaCl, 1 mM DTT and increasing concentrations of imidazole following the manufacturer’s recommendations. Eluted proteins were concentrated, and buffer-exchanged to 10 mM Tris-HCl, 100 mM KCl, 1 mM DTT, 0.1 mM EDTA, pH 7.4 at 25°C using Amicon Ultra-4 Centrifugal 30 kDa Filter Units (Sigma Aldrich NV). Glycerol (final concentration: 50% v/v), Tween 20 (0.5%), Nonidet P40 (0.5%) were then added for long term storage at −20°C.

To quantify protein concentration, a series of Bovine Serum Albumin (BSA) standards in the 100 μg/ml to 1,200 μg/ml range were run in parallel to the concentrated proteins in an SDS-PAGE gel to establish a BSA standard curve used to determine the concentration of the mutants. For the quantification of the commercial phi29 DNAP, the ML (Modified Lowry) Protein Assay (G-Biosciences) was used following the Microtube ML-Protein Assay Protocol (1.5-2.0ml Assay Tubes).

### Primer Extension Assay for HNA synthesis

Reactions were prepared by mixing in 10 μl: 1× NEB 10X Phi29 DNAP buffer, 0.1 mg BSA, 1 M Betaine, 0.2 mM hNTPs, 12 nM phi29 DNAP mutant and 10 pmol of single stranded DNA template, TempN-exoR (**Table S2**) pre-annealed to 1 pmol of fluorescently labelled DNA primer, Tag01F3-exoR (**Table S2**). Pre-annealing was carried out by incubating primer and template at 95°C for 5 min and cooling them down to 4°C at a rate of 0.1°C/sec in a thermocycler. The primer extension assays were incubated at 30°C from 1 min to 16 h (depending on the experiment) but always inactivated by incubating it at 65°C for 20 min. Samples were diluted in an equal volume of 2x loading buffer (98% (v/v) formamide, 10 mM ethylenediaminetetraacetic acid (EDTA), 0.02% (w/v) Orange G) and boiled at 95°C for 5 min. Samples were loaded on 20% TBE-Urea polyacrylamide gels and run in 1X TBE buffer at 25 watts for 2 hours. Gels were imaged using a Typhoon^TM^ FLA 9500 biomolecular imager (GE Healthcare) using recommended filter and detector settings based on the fluorophore being used and analysed with ImageJ (Schneider et al., 2012). Three independent experiments (biological replicates, n = 3) were carried out for the quantification of the primer extension assays.

### DNA Binding Assay – Electrophoretic mobility shift assay (EMSA)

The EMSA protocol was adapted from (Povilaitis et al., 2016). Oligonucleotide Tag01F_ExoR (**Table S2**) was annealed to TemN-ExoR (**Table S2**) at a 1:1.2 molar ratio in the presence of 0.2 M NaCl and 60 mM Tris–HCl, pH 7.5 by incubating at 95°C for 5 min and cooling down to 4°C at a rate of 0.1°C/sec in a thermocycler. A 50 μl complex-forming reaction was prepared by mixing 33 mM Tris-acetate pH 7.9, 66 mM potassium acetate, 10% glycerol, 0.1 mg/ml BSA with the 2 pmol pre-annealed primer/template substrate and corresponding enzyme at a final 60 pM concentration. Reactions were incubated at 30°C for different time intervals (0, 15, 30 or 60 min). After incubation, samples were mixed with loading dye (10% glycerol and 0.03% orange G) and analysed by electrophoresis in 0.8% (w/v) agarose gel in 0.25× TBE buffer (22.25 mM Tris, 22.25 mM boric acid, and 0.5 mM EDTA pH 8.3). Electrophoresis was performed in 0.25x TBE buffer at room temperature for 2 h at 100 V. Gels were imaged using the Typhoon^TM^ bioimager and quantified using ImageQuant TL 8.2 (Cytiva). Four independent experiments (biological replicates, n = 2) were carried out for the quantification of the EMSA assays.

### Rolling Circle Amplification (RCA)

100 pmol of P2_RCA_N8_ExoR (5’-NNNNNN*N*N-3’, **Table S2**) with two phosphorothioated DNA bases at the 3’ end for exonuclease resistance was pre-annealed to 10 ng of pET23_P2_D12A_THR (**Table S1**) plasmid in 1× NEB 10X Phi29 DNAP buffer, per 100 μl RCA reaction, by incubating them at 95°C for 5 min and cooling them down to 4°C at a rate of 0.1°C/sec in a thermocycler. The pre-annealed template mix was supplemented with 0.2 mM dNTPs, 0.2 mg/mL BSA, 5 ng ET SSB (NEB), 4 nM phi29 DNAP in a 100 μl reaction and incubated for 3 h at 30°C. The RCA products were digested with 10U of NotI in 1× Cutsmart buffer for 1 h at 37*C. The digested RCA products were loaded on an agarose gel for visualisation.

### Isothermal polymerase fidelity assay

Single-stranded template was generated by mixing 1.8 pmol of pET23_KOD_DA_Mut (**Table S1**) with 0.7U/μl Nb.BbvCI, 10U/μl of ExoIII and 1× Cutsmart buffer in a 30 μl reaction. 10 pmol of PH_pET23_DA_Biotin3 (**Table S2**) biotinylated oligonucleotide (4.9 μl) was pre-annealed to 0.2 pmol of single stranded template by incubating the mixture at 95°C for 5 min and cooling them down to 4°C at a rate of 0.1°C/sec in a thermocycler. The annealed primer-template was made up to 50 μl with deionized distilled water (ddH_2_O). 5 μl of Dyanabeads MyOne C1 (ThermoFisher), per reaction, were washed twice with 100 μl of 2× BWBS-T (20 mM Tris-HCl pH 7.4, 2 M NaCl, 0.2% v/v Tween20, 2 mM EDTA) and blocked in 1 mL 2× BWBS-T on a rotator for 1 h at room temperature. Beads were captured, resuspended in 50 μl 2x BWBS-T, and mixed with the 50 μl reaction prior to being incubated for 3 h on a rotator at room temperature. Beads were washed twice with 200 μl 30 mM NaOH at 37°C, then with 200 μl EB-T (10 mM Tris-HCl pH 8.8, 0.1 mM EDTA, 0.01% Tween20). Beads were resuspended in 10 μl elution buffer (10 mM Tris-HCl, pH 8.5). Amplicons for deep sequencing were generated using KOD Xtreme™ Hot Start DNA Polymerase (EMD Millipore) following the manufacturer’s instructions in 50 μl reactions with 1 μL of the resuspended beads as template, and outnest_1 (**Table S2**) and P2_fidelity_inestR1 (**Table S2**) as primers. The cycling parameters used were 2 min at 95°C, followed by 20 cycles of 15 s at 98°C, 15 s at 65°C and 10 s at 68°C, with a final polishing step of 2 min at 72°C. Primers were degraded by adding 0.8U/ul Exonuclease I to each 50 μl PCR reaction.

### Next Generation Sequencing of fidelity assay data and analysis

The amplicons described above were purified with the Monarch PCR and DNA purification kit (New England Biolabs) and sent for NGS EZ Amplicon Sequencing by Genewiz. NGS data was pre-processed in the Galaxy public server (usegalaxy.org) using the *NGS_preprocessing.ga* script (**Supplementary information S2**) and trimmed using part 1 of the Julia (v1.7) *Fidelity_Quantification.jl* script (**Supplementary Information S5**). The reference sequence, ‘Fidelity_ref’ (**Table S2**), was added to the trimmed reads and the reads were then aligned using the random chain algorithm of MAFFT version 7 (Katoh & Standley, 2013) on the MAFFT server (mafft.cbrc.jp/alignment/server/large.html) with the following parameters: *mafft --thread 8 --threadtb 5 --threadit 0 --inputorder --randomchain input > output*. The number of insertions, deletions, and substitutions as well as the number of individual error types were calculated by comparing in a per-base manner the aligned reads to the reference ‘Fidelity_ref’ (**Table S2**) sequence within the alignment using part 2 of the *Fidelity_Quantification.jl* script (**Supplementary Information S5**). Raw error rates were obtained by dividing the sum of the errors by the total number of bases sequenced. The PCR and NGS error (R) estimate for Exo+THR was obtained by subtracting the expected number of errors of the commercial phi29 DNAP (using the published error rate 9.5×10^−6^) from the observed number of errors by Exo+THR as follows:

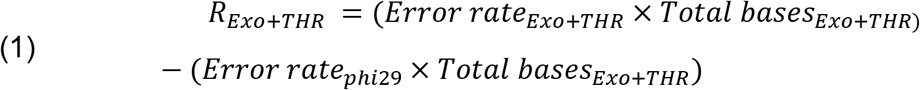

The *Err_Exo+THR_* score was then used to calculate the PCR and NGS error of the D12A-THR and p562del mutants as follows:

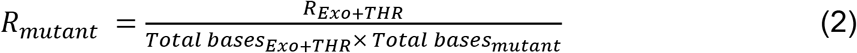

The corrected total error rates were then calculated by dividing the corrected error number over the total bases sequenced from each variant.

Entropy scores were calculated using part 2 of the *Fidelity_Quantification.jl* script (**Supplementary Information S4**) using the following equation:

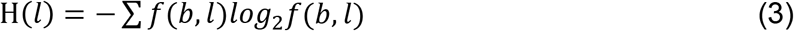

where H(*l*) is the entropy at position *l, b* is a residue or gap and *f*(*b, l*) is the frequency at which residue *b* is found at position *l*.

The location (start position, end position and length) and frequency of each Indel across the MSA were obtained using part2 of the *Fidelity_Quantification.jl* script (**Supplementary Information S5**), by comparing each sequence to the ‘Fidelity_ref’ sequence within the alignment. The frequency values were divided by the total number of insertions or deletions and multiplied by 100 to obtain a relative frequency distribution. Only sequences >5% relative frequency were plotted.

## AVAILABILITY

All raw and processed data used in this manuscript are available in our GitHub repository (https://github.com/PinheiroLab/).

## ACCESSION NUMBERS

Next generation sequencing data has been deposited on NCBI SRA under the following accession number: PRJNA883233.

## SUPPLEMENTARY DATA

SI_Phi29_HNA_synthesis.docx (**Supplementary Information**)

NGS_preprocessing.ga script (**Supplementary Information S2**)

InDel_Quantification.jl (**Supplementary Information S3**)

InDel_EC.xlsx (**Supplementary Information S4**)

Fidelity_Quantification.jl (**Supplementary Information S5**)

## ACKNOWLEDGEMENTS

VBP, LLT and PHM thank ERASynBio (grant BB/N01023X/1; *invivo*XNA). VBP and PHM thank FWO (grant G0H7618N). PHM thanks FWO (studentship 3M180645).

## CONTRIBUTIONS

VBP, LLT and PHM contributed to the project design. PHM performed all experimental work. VBP and PHM carried out NGS analysis.

