## Supplementary_information for "Directed evolution of mesophilic HNA polymerases providing insight into DNA polymerase mechanisms"

**Authors and affiliations:**

Paola Handal-Marquez[1,2], Leticia L. Torres[1] and Vitor B. Pinheiro[2*]

[1] University College London, Department of Structural and Molecular Biology, Gower Street, WC1E 6BT, London, UK

[2] KU Leuven, Rega Institute for Medical Research, Department of Pharmaceutical and Pharmacological Sciences, Herestraat, 49 – box 1041, 3000 Leuven, Belgium

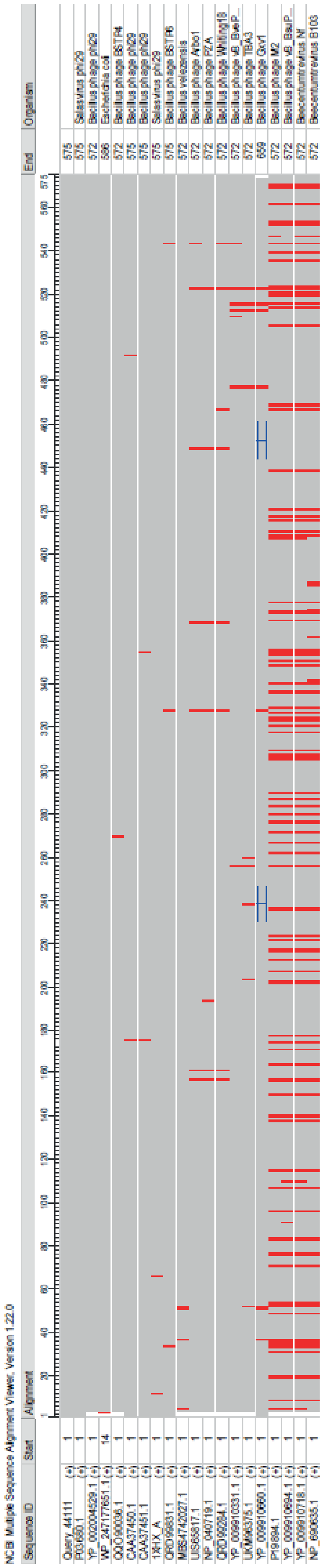

**Figure S1. Phi29 DNAP homologues in public databases.** Sequences with >80% sequence identity and >80% query cover from a blast search of the Phi29 DNAP protein sequence were selected, aligned using the NCBI Multiple Alignment Tool (Altschul et al., 1990) and viewed using the NCBI'S Sequence Viewer (Rangwala et al., 2021). Mismatches relative to the query (Phi29 DNAP sequence) are shown in red and insertions are indicated by a blue bracket. Only 20 sequences show significant similarity to phi29 DNAP.

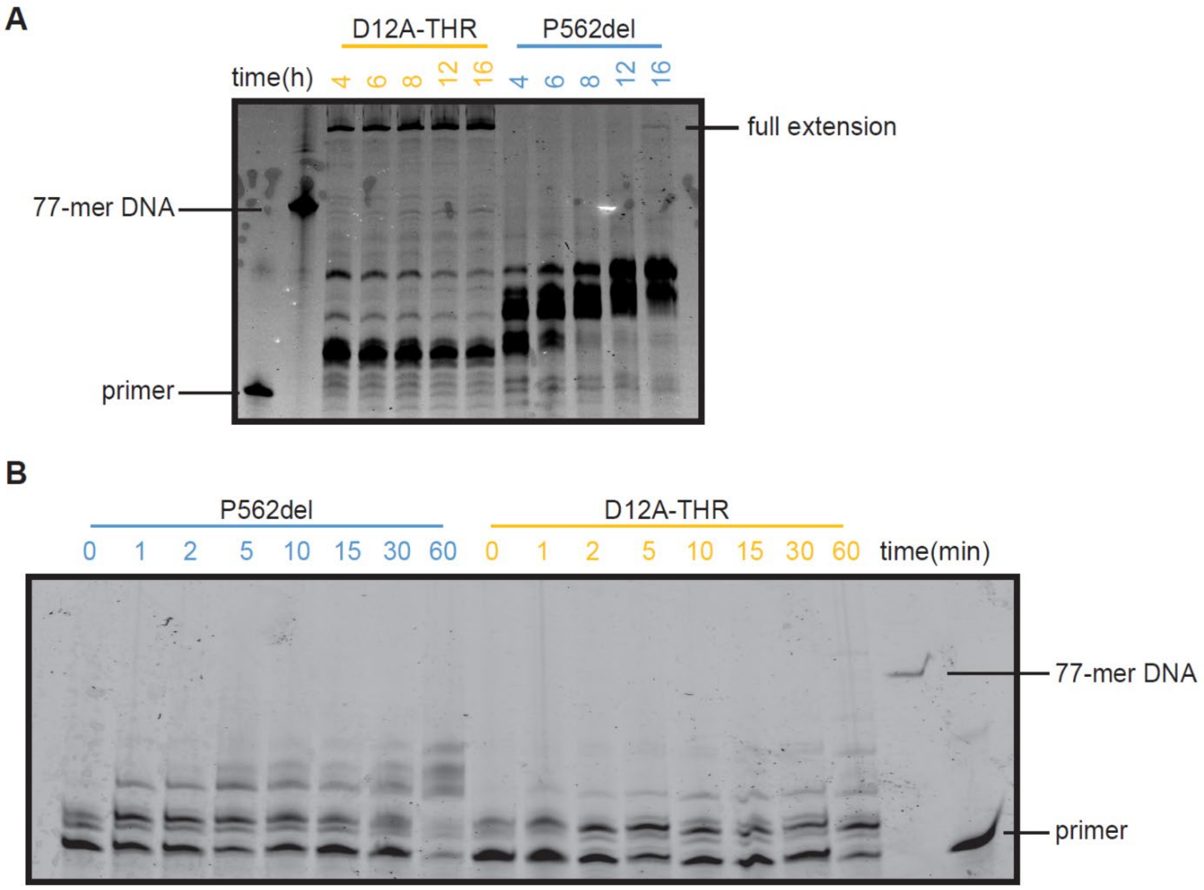

**Figure S2. HNA synthesis time courses by D12A-THR and p562del with different templates.** Products from primer extensions by D12A-THR and P562 mutants on the TempN-exoR (Table S2) template with incubation times from 4 to 16 h (**A**) as well as on the TempN_2.7_ExoR template (Table S2) with incubation times from 0 to 1 h (**B**) were separated by denaturing PAGE. The TempN_2.7_ExoR template is a modified version of TempN-exoR with 4 substitutions and 1 deletion that reduces the probability of secondary structure formation. Fully extended products (57 hNTP incorporations) are shown. HNA migrates slower than DNA in denaturing PAGE (Torres & Pinheiro, 2018).

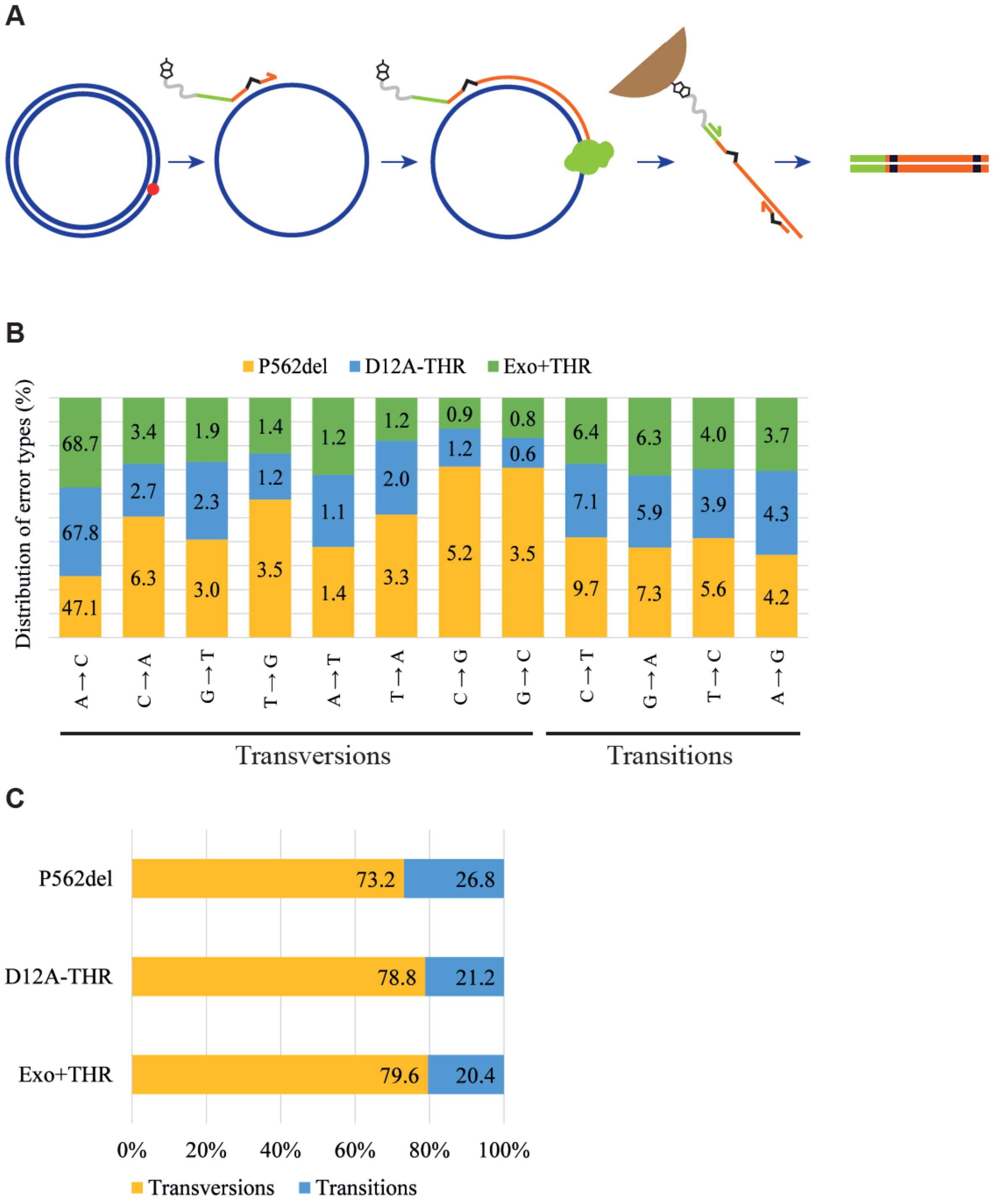

**Figure S3. Phi29 DNAP P562del reduced fidelity and increased InDel incorporations rate.** (**A**) Workflow of the isothermal polymerase fidelity assay. The red dot marks the nicking site for single stranded plasmid generation, the non-complementary overhang of the primer for downstream amplification is shown in green and 1 bp mismatches are shown in black. The primer is extended, captured, and purified through biotin-streptavidin pulldown and used as template in a secondary PCR amplification step. (**B**) Distribution and quantification of error (misincorporation) types introduced by each mutant during isothermal DNA replication. Each error type was identified by comparing the isothermal amplification products post-deep sequencing and after their alignment to the Fidelity_ref (Table S2) on a base-to-base manner. The sum of each error type was divided by the total number of misincorporations and multiplied times a 100 to yield the error type percentage displayed. (**C**) The total percentage of inversions and transversions introduced by each mutant.

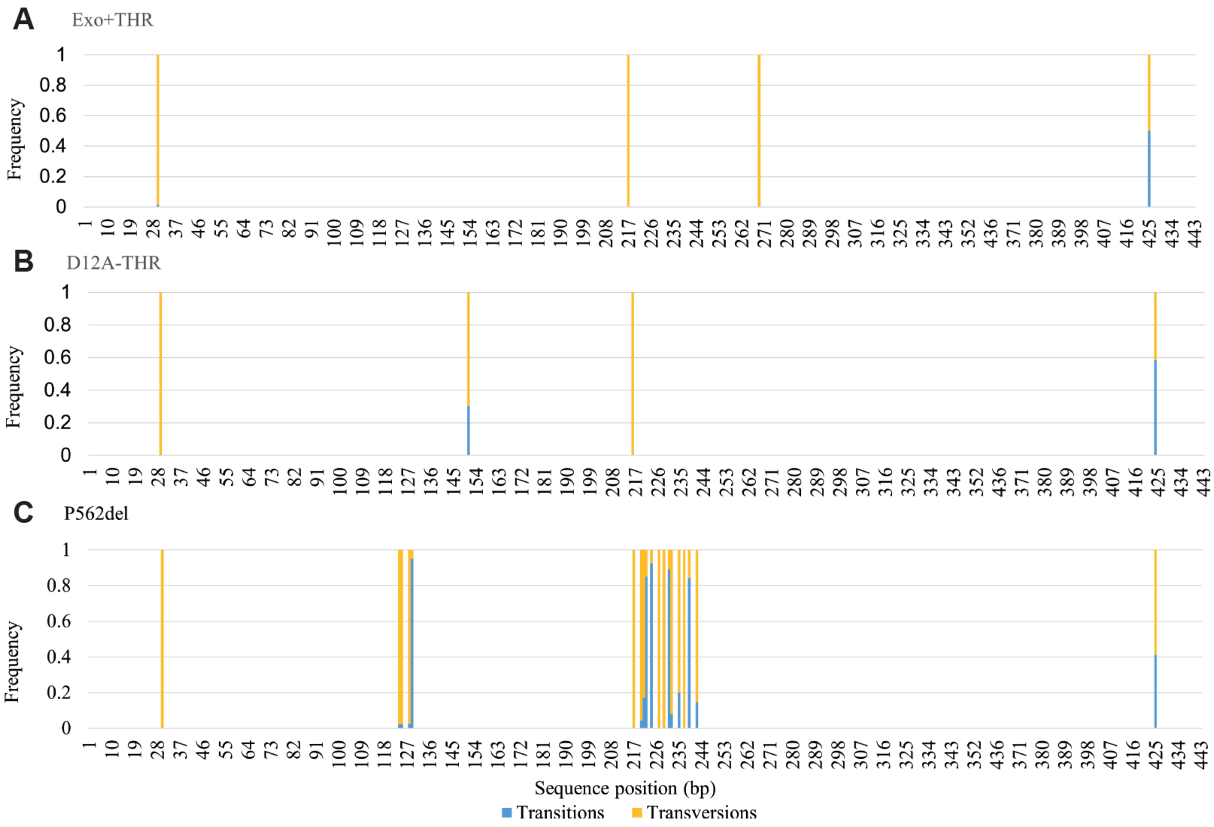

**Figure S4. Transition and transversion hotspots introduced by phi29 DNAP variants.** The products from the isothermal DNA replication fidelity assays generated by Exo+THR, D12A-THR and p562del were deep sequenced, filtered by quality, trimmed, and aligned. The MSA alignments were used to quantify the abundance of transitions and transversions per position by comparing each of the aligned reads to the Fidelity_ref (table S2) sequence within each the alignment in a base-by-base manner. The total number of transitions and transversions per position was divided by the number of reads to obtain overall frequency scores. Only positions with transitions or transversions with overall frequency scores above 0.5% were selected for visualization. The scores of each error type were divided by the sum of both scores to obtain the frequency value per position and were plotted against the Fidelity_ref length.

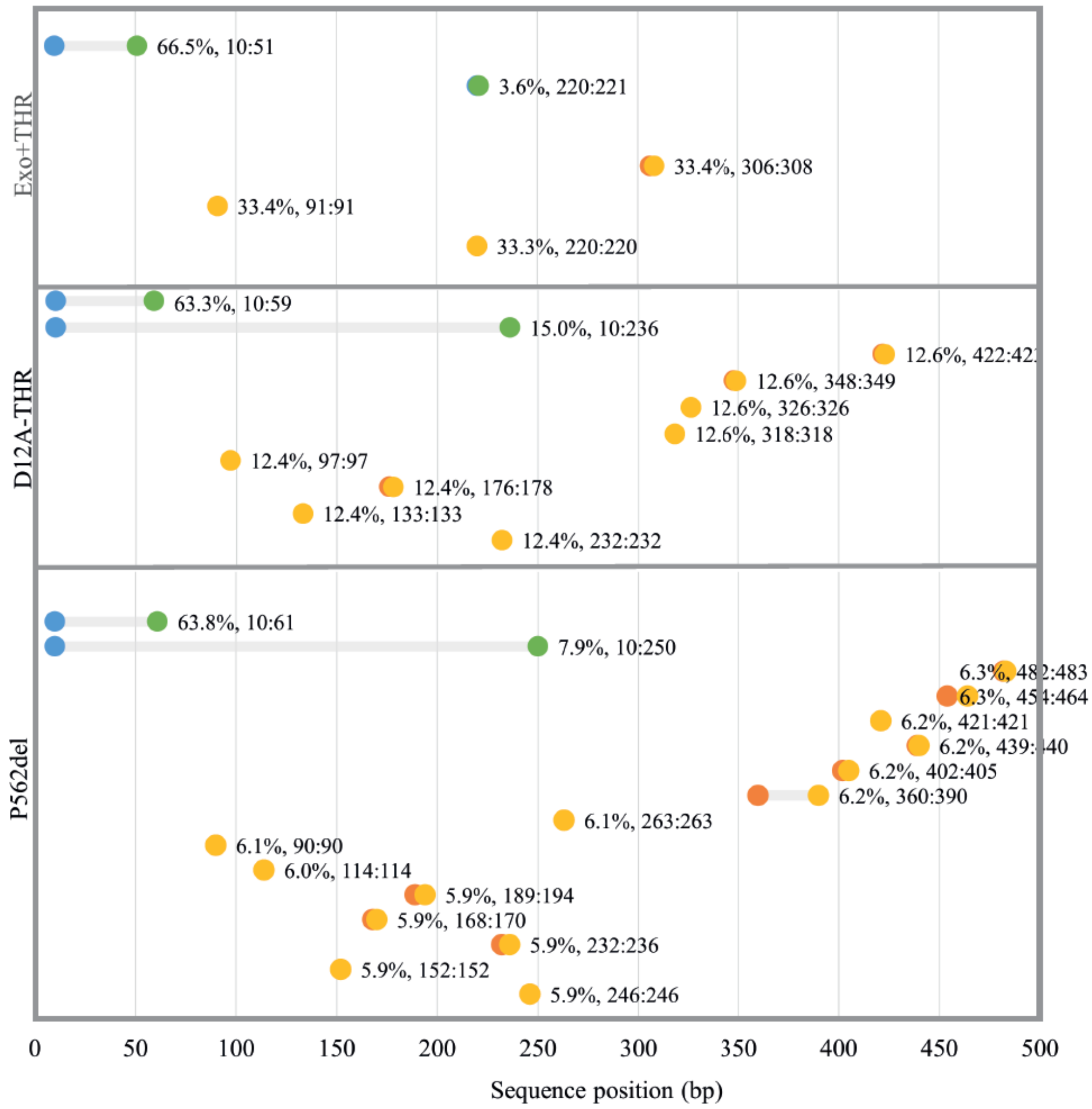

**Figure S5. Location of most abundant InDel introductions by each mutant during isothermal DNA replication.** Location of deletions (blue to green dots) and insertions (orange to yellow dots) appearing with >5% frequency relative to all the insertions or deletions identified in the MSA of the isothermal amplification products generated by each mutant. The x-axis indicates the sequence length of the template used in the assay/analysis. Blue or orange dots indicate the 'start' of the deletion or insertion respectively, and the green or yellow dots indicate the 'end' of the deletion or insertion respectively. The percentage values adjacent to the ‘end’ dots represent the abundance of deletions or insertions relative to the total number of deletions or insertions respectively. The percentage values are followed the location of the particular InDel in a range format.

**Table S1. Sequences[a] of all the plasmids used in this study.**

| pET23-P2-D12A-THR |
|---|
| acgcgccctgtagcggcgcattaagcgcggccgctgtggtggttacgcgcagcgtgaccgctacacttgccagcgcc |
| ctagcgcccgctcctttcgctttcttcccttcctttctcgccacgttcgccggctttccccgtcaagctctaaatcgggggctcc |
| ctttagggttccgatttagtgctttacggcacctcgaccccaaaaaacttgattagggtgatggttcacgtagtgggccatc |
| gccctgatagacggtttttcgccctttgacgttggagtccacgttctttaatagtggactcttgttccaaactggaacaacact |
| caaccctatctcggtctattcttttgatttataagggattttgccgatttcggcctattggttaaaaaatgagctgatttaacaa |
| aaatttaacgcgaattttaacaaaatattaacgtttacaatttcaggtggcacttttcggggaaatgtgcgcggaacccct |
| atttgtttatttttctaaatacattcaaatatgtatccgctcatgagacaataaccctgataaatgcttcaataatattgaaaaa |
| ggaagagtatgagtattcaacatttccgtgtcgcccttattcccttttttgcggcattttgccttcctgtttttgctcacccagaaa |
| cgctggtgaaagtaaaagatgctgaagatcagttgggtgcacgagtgggttacatcgaactggatctcaacagcggta |
| agatccttgagagttttcgccccgaagaacgttttccaatgatgagcacttttaaagttctgctatgtggcgcggtattatcc |
| cgtattgacgccgggcaagagcaactcggtcgccgcatacactattctcagaatgacttggttgagtactcaccagtca |
| cagaaaagcatcttacggatggcatgacagtaagagaattatgcagtgctgccataaccatgagtgataacactgcg |
| gccaacttacttctgacaacgatcggaggaccgaaggagctaaccgcttttttgcacaacatgggggatcatgtaactc |
| gccttgatcgttgggaaccggagctgaatgaagccataccaaacgacgagcgtgacaccacgatgcctgcagcaat |
| ggcaacaacgttgcgcaaactattaactggcgaactacttactctagcttcccggcaacaattaatagactggatggag |
| gcggataaagttgcaggaccacttctgcgctcggcccttccggctggctggtttattgctgataaatctggagccggtga |
| gcgtgggtctcgcggtatcattgcagcactggggccagatggtaagccctcccgtatcgtagttatctacacgacgggg |
| agtcaggcaactatggatgaacgaaatagacagatcgctgagataggtgcctcactgattaagcattggtaactgtca |
| gaccaagtttactcatatatactttagattgatttaaaacttcatttttaatttaaaaggatctaggtgaagatcctttttgataat |
| ctcatgaccaaaatcccttaacgtgagttttcgttccactgagcgtcagaccccgtagaaaagatcaaaggatcttcttga |
| gatcctttttttctgcgcgtaatctgctgcttgcaaacaaaaaaaccaccgctaccagcggtggtttgtttgccggatcaag |
| agctaccaactctttttccgaaggtaactggcttcagcagagcgcagataccaaatactgtacttctagtgtagccgtagt |
| taggccaccacttcaagaactctgtagcaccgcctacatacctcgctctgctaatcctgttaccagtggctgctgccagtg |
| gcgataagtcgtgtcttaccgggttggactcaagacgatagttaccggataaggcgcagcggtcgggctgaacgggg |
| ggttcgtgcacacagcccagcttggagcgaacgacctacaccgaactgagatacctacagcgtgagctatgagaaa |
| gcgccacgcttcccgaagggagaaaggcggacaggtatccggtaagcggcagggtcggaacaggagagcgcac |
| gagggagcttccagggggaaacgcctggtatctttatagtcctgtcgggtttcgccacctctgacttgagcgtcgatttttgt |

gatgctcgtcaggggggcggagcctatggaaaaacgccagcaacgcggcctttttacggttcctggccttttgctgcgtt
atcccctgattctgtggccttttgcgcgtctgcgttatcccctgattctgatgttctttcctgcgttatcccctgattctgtggataa
ccgtattaccgcctttgagtgagctgcgttatcccctgattctgctgataccgctcgccgcagccgaacgaccgagcgca
gcgagtcagtgagcgaggaagcggaatatcgcctgatgcggtattttctccttacgcatctgtgcggtatttcacaccgc
aatggtgcactctcagtacaatctgctctgatgccgcatagttaagccagtatacactccgctatcgctacgtgactgggt
catggctgcgccccgacacccgccaacacccgctgacgcgccctgacgggcttgtctgctcccggcatccgcttaca
gacaagctgtgaccgtctccgggagctgcatgtgtcagaggttttcaccgtcatcaccgaaacgcgcgaggcagctgc
ggtaaagctcatcagcgtggtcgtgaagcgattcacagatgtctgcctgttcatccgcgtccagctcgttgagtttctccag
aagcgttaatgtctggcttctgataaagcgggccatgttaagggcggtttttcctgtttggtcactgatgcctccgtgtaagg
gggatttctgttcatgggggtaatgataccgatgaaacgagagaggatgctcacgatacgggttactgatgatgaacat
gcccggttactggaacgttgtgagggtaaacaactggcggtatggatgcggcgggaccagagaaaaatcactcagg
gtcaatgccagcgcttcgttaatacagatgtaggtgttccacagggtagccagcagcatcctgcgatgcagatccgga
acataatggtgcagggcgctgacttccgcgtttccagactttacgaaacacggaaaccgaagaccattcatgttgttgct
caggtcgcagacgttttgcagcagcagtcgcttcacgttcgctcgcgtatcggtgattcattctgctaaccagtaaggcaa
ccccgccagcctagccgggtcctcaacgacaggagcacgatcatgcgcacccgtggccaggacccaacgctgccc
gagatctcgatcccgcgaaattaatacgactcactatagggagaccacaacggtttccctctagaaataattttgtttaac
tttaagaaggagatataccatggatcctctagagtcgacctgcaggcatgcaagcttgcggccacacaggagatagtc
atacatgaaacacatgcctcgcaaaaggtatagctgcgcttttgaaaccaccaccaaagttgaagattgtcgtgtttggg
catatggctatatgaacattgaagatcacagcgagtataaaatcggcaatagcctggatgaatttatggcatgggctctg
aaagttcaggccgatctgtattttcacaatctgaaatttgatggtgccttcattattaactggctggaacgcaatggttttaaa
tggtcagcagatggtctgccgaatacctataacaccattattagccgtacgggccagtggtatatgattgatatttgcctgg
gttataaaggcaaacgcaaaattcataccgtgatctatgacagcctgaaaaaactgccgtttccggtgaaaaaaatcg
ccaaagatttcaaactgaccgtgctgaaaggcgatatcgattatcacaaagaacgtccggttggctacaaaattacac
cggaagaatatgcctacatcaaaaacgacattcagattattgcagaagccctgctgattcagtttaaacagggtctggat
cgtatgaccgcaggtagcgatagcctgaaagattttaaagatatcattaccaccaaaaaattcaaaaaagtgttcccga
ccctgagcctgggcctggataaaaaagttcgttacgcatatcgcggtggttttacctggctgaatgatcgctttaaagaaa
aagaaattggcgagggcatggtgtttgatgttaatagcctgtatccggcacagatgtatagccgtctgctgccgtatggtg
aaccgattgtttttgaaggtaaatatgtgtgggatgaggattatccgctgcatattcagcatattcgttgcgaatttgaactga
aagaaggctatattccgaccattcagatcaaacgtagccgcttctataaaggtaacgagtatctgaaaagcagcggtg
gtgaaattgcagatctgtggctgagcaatgttgatctggaactgatgaaagaacactacgatctgtacaacgtggaatat
atcagcggtctgaaattcaaagcaaccaccggtctgttcaaagacttcattgataaatggacctatatcaaaaccacctc

cgaaggtgcaattaaacagctggcaaaactgatgctgaattccctgtatggtaaatttgcaagcaatccggatgtgacc
ggtaaagttccttatctgaaagaaaatggtgcactgggttttcgtctgggtgaagaagaaaccaaagatccggtttatac
cccgatgggtgtgtttattaccgcatgggcacgttataccaccattaccgcagcacaggcatgttatgaccgtattatctatt
gtgataccgatagcattcatctgaccggcaccgaaattccggatgttatcaaagatattgtggatcctaaaaaactgggc
tattgggcacatgaaagcacctttaaacgtgcaaaatatctgcgcccgaaaacctatatccaggatatctatatgaaag
aagtggacggtgaactggttgcaggtagtccggatgattacaccgatatcaaacttagcgttaaatgtgccggtatgac
cgataaaatcaaaaaagaagtgaccttcgagaacttcaaagtgggttttagccgtaaaatgaaaccgaaaccggttc
aggttccgggtggtgttgttctggttgatgatacctttacgatcaaacaccaccaccaccaccactgagatccggctgcta
acaaagcccgaaaggaagctgagttggctgctgccaccgctgagcaataactagcataaccccttggggcctctaaa
cgggtcttgaggggttttttgctgaaaggaggaactatatccggattggcgaatggg

**pET23_KOD_DA_Mut**

tggcgaatgggacgcgccctgtagcggcgcattaagcgcggccgctgtggtggttacgcgcagcgtgaccgctacac
ttgccagcgccctagcgcccgctcctttcgctttcttcccttcctttctcgccacgttcgccggctttccccgtcaagctctaaa
tcgggggctccctttagggttccgatttagtgctttacggcacctcgaccccaaaaaacttgattagggtgatggttcacgt
agtgggccatcgccctgatagacggtttttcgccctttgacgttggagtccacgttctttaatagtggactcttgttccaaact
ggaacaacactcaaccctatctcggtctattcttttgatttataagggattttgccgatttcggcctattggttaaaaaatgag
ctgatttaacaaaaatttaacgcgaattttaacaaaatattaacgtttacaatttcaggtggcacttttcggggaaatgtgcg
cggaacccctatttgtttatttttctaaatacattcaaatatgtatccgctcatgagacaataaccctgataaatgcttcaata
atattgaaaaaggaagagtatgagtattcaacatttccgtgtcgcccttattcccttttttgcggcattttgccttcctgtttttgct
cacccagaaacgctggtgaaagtaaaagatgctgaagatcagttgggtgcacgagtgggttacatcgaactggatct
caacagcggtaagatccttgagagttttcgccccgaagaacgttttccaatgatgagcacttttaaagttctgctatgtggc
gcggtattatcccgtattgacgccgggcaagagcaactcggtcgccgcatacactattctcagaatgacttggttgagta
ctcaccagtcacagaaaagcatcttacggatggcatgacagtaagagaattatgcagtgctgccataaccatgagtga
taacactgcggccaacttacttctgacaacgatcggaggaccgaaggagctaaccgcttttttgcacaacatgggga
tcatgtaactcgccttgatcgttgggaaccggagctgaatgaagccataccaaacgacgagcgtgacaccacgatgc
ctgcagcaatggcaacaacgttgcgcaaactattaactggcgaactacttactctagcttcccggcaacaattaataga
ctggatggaggcggataaagttgcaggaccacttctgcgctcggcccttccggctggctggtttattgctgataaatctgg
agccggtgagcgtgggtctcgcggtatcattgcagcactggggccagatggtaagccctcccgtatcgtagttatctaca
cgacggggagtcaggcaactatggatgaacgaaatagacagatcgctgagataggtgcctcactgattaagcattgg
taactgtcagaccaagtttactcatatatactttagattgatttaaaacttcatttttaatttaaaaggatctaggtgaagatcct

ttttgataatctcatgaccaaaatcccttaacgtgagttttcgttccactgagcgtcagaccccgtagaaaagatcaaagg
atcttcttgagatcctttttttctgcgcgtaatctgctgcttgcaaacaaaaaaaccaccgctaccagcggtggtttgtttgcc
ggatcaagagctaccaactctttttccgaaggtaactggcttcagcagagcgcagataccaaatactgtacttctagtgt
agccgtagttaggccaccacttcaagaactctgtagcaccgcctacatacctcgctctgctaatcctgttaccagtggctg
ctgccagtggcgataagtcgtgtcttaccgggttggactcaagacgatagttaccggataaggcgcagcggtcgggct
gaacggggggttcgtgcacacagcccagcttggagcgaacgacctacaccgaactgagatacctacagcgtgagct
atgagaaagcgccacgcttcccgaagggagaaaggcggacaggtatccggtaagcggcagggtcggaacagga
gagcgcacgagggagcttccagggggaaacgcctggtatctttatagtcctgtcgggtttcgccacctctgacttgagcg
tcgatttttgtgatgctcgtcaggggggcggagcctatggaaaaacgccagcaacgcggcctttttacggttcctggcctt
ttgctgcgttatcccctgattctgtggccttttgcgcgtctgcgttatcccctgattctgatgttctttcctgcgttatcccctgattct
gtggataaccgtattaccgcctttgagtgagctgcgttatcccctgattctgctgataccgctcgccgcagccgaacgacc
gagcgcagcgagtcagtgagcgaggaagcggaatatcgcctgatgcggtattttctccttacgcatctgtgcggtatttc
acaccgcaatggtgcactctcagtacaatctgctctgatgccgcatagttaagccagtatacactccgctatcgctacgt
gactgggtcatggctgcgccccgacacccgccaacacccgctgacgcgccctgacgggcttgtctgctcccggcatc
cgcttacagacaagctgtgaccgtctccgggagctgcatgtgtcagaggttttcaccgtcatcaccgaaacgcgcgag
gcagctgcggtaaagctcatcagcgtggtcgtgaagcgattcacagatgtctgcctgttcatccgcgtccagctcgttga
gtttctccagaagcgttaatgtctggcttctgataaagcgggccatgttaagggcggttttttcctgtttggtcactgatgcctc
cgtgtaagggggatttctgttcatgggggtaatgataccgatgaaacgagagaggatgctcacgatacgggttactgat
gatgaacatgcccggttactggaacgttgtgagggtaaacaactggcggtatggatgcggcgggaccagagaaaaa
tcactcagggtcaatgccagcgcttcgttaatacagatgtaggtgttccacagggtagccagcagcatatggtgcaggg
cgctgacttccgcgtttccagactttacgaaacacggaaaccgaagaccattcatgttgttgctcaggtcgcagacgtttt
gcagcagcagtcgcttcacgttcgctcgcgtatcggtgattcattctgctaaccagtaaggcaaccccgccagcctagc
cgggtcctcaacgacaggagcacgatcatgcgcacccgtggccaggacccaacgctgcccgagatctcgatcccg
cgaaattaatacgactcactatagggagaccacaacggtttccctctagaaataattttgtttaactttaagaaggagata
taccatggatcctctagagtcgacctgcaggcatgcaagcttgcggccacacaggagatagtcatacatgaaacaca
aagaggagaaattaactatgagaggatctcaccatcaccatcaccatacggatccaagcggcctggtgccgcgcgg
cagcatgatcctcgacactgactacataaccgaggatggaaagcctgtcataagaattttcaagaaggaaaacggcg
agtttaagattgagtacgaccggacttttgaaccctacttctacgccctcctgaaggacgattctgccattgaggaagtca
agaagataaccgccgagaggcacgggacggttgtaacggttaagcgggttgaaaaggttcagaagaagttcctagg
gagaccagttgaggtctggaaactctactttactcatccgcaggacgaaccagcgataagggacaagatacgagag
catccagcagttattgacatctacgagtacgacatacccttcgccaagcgctacctcatagacaagggattagtgccaa

| |
|---|
| tggaaggcgacgaggagctgaaaatgctcgccttcgcgattgcgactctctaccatgagggcgaggagttcgccgag<br>gggccaatccttatgataagctacgccgacgaggaaggggccagggtgataacttggaagaacgtggatctcccta<br>cgttgacgtcgtctcgacggagagggagatgataaagcgcttcctccgtgttgtgaaggagaaagacccggacgttct<br>cataacctacaacggcgacaacttcgacttcgcctatctgaaaaagcgctgtgaaaagctcggaataaacttcgccct<br>cggaagggatggaagcgagccgaagattcagaggatgggcgacaggtttgccgtcgaagtgaagggacggatac<br>acttcgatctctatcctgtgataagacggacgataaacctgcccacatacacgcttgaggccgtttatgaagccgtcttcg<br>gtcagccgaaggagaaggtttacgctgaggaaataaccacagcctgggaaaccggcgagaaccttgagagagtcg<br>cccgctactcgatggaagatgcgaaggtcacatacgagcttgggaaggagttccttccgatggaggcccagctttctcg<br>cttaatcggccagtccctctgggacgtctcccgctccagcactggcaacctcgttgagtggttcctcctcaggaaggcct<br>atgagaggaatgagctggccccgaacaagcccgatgaaaaggagctggccagaagacggcagagctatgaagg<br>aggctatgtaaaagagcccgagagagggttgtgggagaacatagtgtacctagattttagatccctgtacccctcaatc<br>atcatcacccacaacgtctcgccggatacgctcaacagagaaggatgcaaggaatatgacgttgccccacaggtcg<br>gccaccgcttctgcaaggacttcccaggatttatcccgagcctgctaggagacctcctagaggagaggcagaagata<br>aagaagaagatgaaggccacgattgacccgatcgagaggaagctcctcgattacaggcagaggttgatcaagatcc<br>tggcaaacagctactacggttactacggctatgcaagggcgcgctggtactgcaaggagtgtgcagagagcgtaacg<br>gcctggggaagggagtacataacgatgaccatcaaggagatagaggaaaagtacggctttaaggtaatctacagcg<br>acaccgacggatttttgccacaatacctggagccgatgctgaaaccgtcaaaaagaaggctatggagttcctcaagt<br>atatcaacgccaaacttccgggcgcgcttgagctcgagtacgagggcttctacaaacgcggcttcttcgtcacgaaga<br>agaagtatgcggtgatagacgaggaaggcaagataacaacgcgcggacttgagattgtgaggcgtgactggagcg<br>agatagcgaaagagacgcaggcgagggttcttgaagctttgctaaaggacggtgacgtcgagaaggccgtgaggat<br>agtcaaagaagttaccgaaaagctgagcaagtacgaggttccgccggagaagctggtgatccacgagcagataac<br>gagggatttaaaggactacaaggcaaccggtccccacgttgccgttgccaagaggttggccgcgagaggagtcaaa<br>atacgccctggaacggtgataagctacatcgtgctcaagggctctgggaggataggcgacagggcgataccgttcga<br>cgagttcgacccgacgaagcacaagtacgacgccgagtactacattgagaaccaggttctcccagccgttgagaga<br>attctgagagccttcggttaccgcaaggaagacctgcgctaccagaagacgagacaggttggtttgagtgcttggctga<br>agccgaagggaacttgatcgatgctccgagatgaggtaggatggctggcttacggtgttactaatgagccatcctcgag<br>caccaccaccaccaccactgagatccggctgctaacaaagcccgaaaggaagctgagttggctgctgccaccgctg<br>agcaataactagcataaccccttggggcctctaaacgggtcttgaggggttttttgctgaaaggaggaactatatccgga<br>t |

[a]All sequences are written in the 5’→3’ direction

**Table S2. Sequences[a] of the oligonucleotides and templates used in this study.**

| Category | Name | Sequence[b] |
|---|---|---|
| Exo loop InDel mutagenesis | Exo_loop_R | TTCAACTTTGGTGGTGGTTTC |
| | Exo_loop_INS1 | NNSGATTGTCGTGTTTGGGCATATG |
| | Exo_loop_INS2 | NNSNNSGATTGTCGTGTTTGGGCATATG |
| | Exo_loop_INS3 | NNSNNSNNSGATTGTCGTGTTTGGG |
| | Exo_loop_DEL1 | TGTCGTGTTTGGGCATATGG |
| | Exo_loop_DEL2 | CGTGTTTGGGCATATGGCTATATG |
| | Exo_loop_DEL3 | GTTTGGGCATATGGCTATATGAAC |
| TPR2 loop InDel mutagenesis | TPR2_loop_R | TTCTTTCAGATAAGGAACTTTACC |
| | TPR2_loop_INS2 | NNSNNSAATGGTGCACTGGGT |
| | TPR2_loop_INS1 | NNSAATGGTGCACTGGG |
| | TPR2_loop_INS3 | NNSNNSNNSAATGGTGCACTGGG |
| | TPR2_loop_DEL1 | GGTGCACTGGGTTTTC |
| | TPR2_loop_DEL2 | GCACTGGGTTTTCGTC |
| Thumb loop InDel mutagenesis | Thumb_loop_R | AACCTGAACCGGTTTCGGTTTC |
| | Thumb_loop_INS2 | NNSNNSCCGGGTGGTGTTGTTC |
| | Thumb_loop_INS1 | NNSCCGGGTGGTGTTGTTCTG |
| | Thumb_loop_INS3 | NNSNNSNNSCCGGGTGGTGTTGTTC |

| | | |
|---|---|---|
| | Thumb_loop_DEL1 | GGTGGTGTTGTTCTGGTTGATGATAC |
| | Thumb_loop_DEL2 | GGTGTTGTTCTGGTTGATGATACCTTTAC |
| | Thumb_loop_DEL3 | GTTGTTCTGGTTGATGATACCTTTACGATC |
| | Thumb_loop_DEL4 | GTTCTGGTTGATGATACCTTTACGATCAAA |
| CST selection | CST_04(7)exoR | /5BiotinTEG/ACC*G*C*A |
| P562del mutant | p2_thumb_loop_R | AACCTGAACCGGTTTCGGTTTC |
| | p2_thumb_loop_DEL1 | GGTGGTGTTGTTCTGGTTGATGATAC |
| Exo+THR mutant | iPCR_P2_Exo+_F1 | GTATAGCTGCGATTTTGAAACC |
| | iPCR_P2_Exo+_R1 | CTTTTGCGAGGCATGTG |
| Primer extension assay | TempN-exoR | TGGTCCAGCATCGTGAGATCGATTACCGAA CAGCACTACGTGGCTAAGTGCTTATCTCCTA GCTTAAACGGAT*C*C*G |
| | TempN_2.7_ExoR | TGGTCCAGCATCGTGAGATCCCTTACTGAA CAGACTACATGGCTAAGTGCTTATCTCCTAG CTTAAACGGAT*C*C*G |
| RCA assay | P2_RCA_N8_ExoR | NNNNNN*N*N |
| Fidelity assay | PH_pET23_DA_Biotin 3 | /5BiotinTEG/CCCCTTATTAGCGTTTGCCAGC TCTTCCACTCAGGGTtAATGCCAGC |
| | outnest_1 | CCCCTTATTAGCGTTTGCCA |
| | P2_fidelity_inestR1 | CTGTGTGGCCGCAAG |
| | Fidelity_ref | agggttaatgccagcgcttcgttaatacagatgtaggtgttccac agggtagccagcagcatatggtgcagggcgctgacttccgcgtt |

| | | |
|---|---|---|
| | | tccagactttacgaaacacggaaaccgaagaccattcatgttgttgctcaggtcgcagacgttttgcagcagcagtcgcttcacgttcgctcgcgtatcggtgattcattctgctaaccagtaaggcaaccccgccagcctagccgggtcctcaacgacaggagcacgatcatgcgcacccgtggccaggacccaacgctgcccgagatctcgatcccgcgaaattaatacgactcactatagggagaccacaacggtttccctctagaaataattttgtttaactttaagaaggagatataccatggatcctctagagtcgacctgcaggcatgcaagcttgcggccacacag |

[a]All sequences are written in the 5’→3’ direction

[b]N: A/C/G/T; S:G/C

*: phosphorothioate bond

/5BiotinTEG/: Biotin with a 15 atom triethylene glycol (TEG) spacer
